# Resolving conformational changes that mediate a two-step catalytic mechanism in a model enzyme

**DOI:** 10.1101/2023.06.02.543507

**Authors:** Jack B. Greisman, Kevin M. Dalton, Dennis E. Brookner, Margaret A. Klureza, Candice J. Sheehan, In-Sik Kim, Robert W. Henning, Silvia Russi, Doeke R. Hekstra

## Abstract

Enzymes catalyze biochemical reactions through precise positioning of substrates, cofactors, and amino acids to modulate the transition-state free energy. However, the role of conformational dynamics remains poorly understood due to lack of experimental access. This shortcoming is evident with *E. coli* dihydrofolate reductase (DHFR), a model system for the role of protein dynamics in catalysis, for which it is unknown how the enzyme regulates the different active site environments required to facilitate proton and hydride transfer. Here, we present ligand-, temperature-, and electric-field-based perturbations during X-ray diffraction experiments that enable identification of coupled conformational changes in DHFR. We identify a global hinge motion and local networks of structural rearrangements that are engaged by substrate protonation to regulate solvent access and promote efficient catalysis. The resulting mechanism shows that DHFR’s two-step catalytic mechanism is guided by a dynamic free energy landscape responsive to the state of the substrate.

## Introduction

Enzymes serve essential cellular functions by selectively enhancing the rates of chemical reactions. This catalysis is often explained through precise positioning of substrates and functional groups to stabilize the transition state of a reaction [1, 2]. Proteins, however, contain many rotatable bonds with energetic barriers that can be crossed by thermal motion. Therefore, proteins exhibit conformational dynamics best described by an ensemble of structures [3, 4]. Since even sub-angstrom changes in important interactions are, in principle, sufficient to impact the energetics of catalytic steps or their allosteric regulation [3, 5–8], conformational changes can be small enough to be overlooked by existing methods, yet key to understanding enzyme function. A central question therefore remains—how do the conformational dynamics of enzymes relate to the chemical reaction coordinate? Better understanding of this relation would have far-reaching implications for the rational design of artificial enzymes, for understanding how function constrains evolution, and in the design of pharmacological modulators of enzyme activity.

Critical gaps in our understanding of the interplay of conformational dynamics and the chemical steps of enzyme catalysis are evident for even the best-studied enzymes. Dihydrofolate reductase (DHFR) from *Escherichia coli* (hereafter, *ec*DHFR) has been studied intensively for decades [9–16]. DHFR catalyzes the stereospecific transfer (Fig. 1A) of a hydride (H*^−^*) from reduced nicotinamide adenine dinucleotide phosphate (NADPH) to dihydrofolate (DHF), yielding NADP^+^ and tetrahydrofolate (THF), an essential precursor for purine synthesis [10]. Kinetic isotope effect measurements support a stepwise catalytic mechanism for *ec*DHFR in which protonation of DHF at the N5 atom precedes hydride transfer [17] (Fig. 1A). A key active-site loop, the Met20 loop, adopts two different conformations depending on the bound ligands: the closed conformation is associated with the Michaelis complex—the catalytically competent state in which the enzyme is bound to its cofactor and substrate, as shown in Fig. 1B. The occluded conformation is, instead, adopted by product complexes to promote exchange of the spent NADP^+^ cofactor [9].

**Fig. 1.**
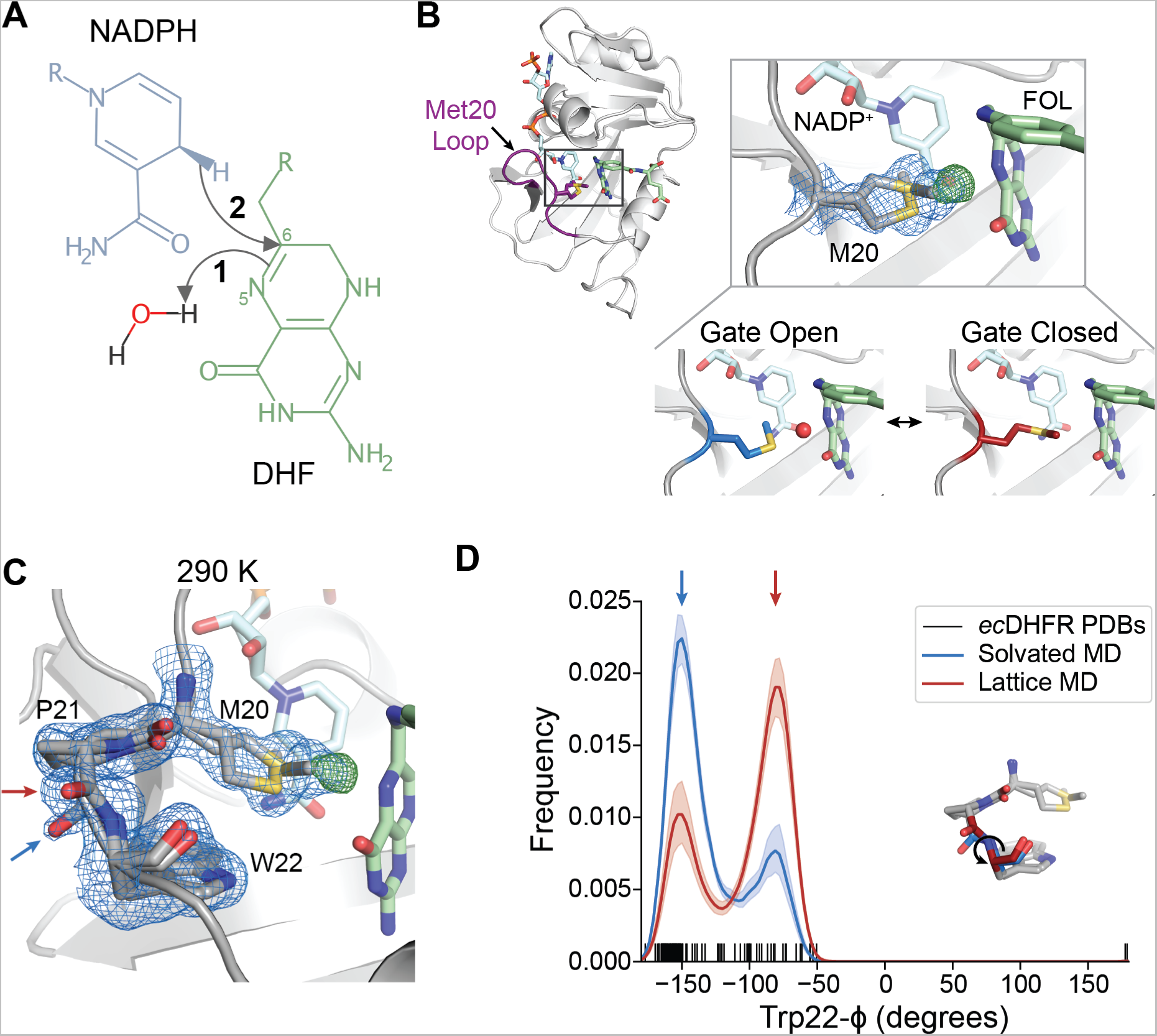
The closed state of the Met20 loop contains two interconverting substates. (A) Schematic of the hydride transfer reaction catalyzed by DHFR. Hydride transfer occurs from NADPH to dihydrofolate (DHF) with a stepwise mechanism: protonation of DHF from water precedes hydride transfer. The N5 nitrogen and C6 carbon of DHF are labeled. (B) and (C) 2*mFo −DFc* map (blue mesh; 0.7*σ*), *mFo −DFc* (green mesh; +4.0*σ*), and refined model for a *ec*DHFR:NADP^+^:FOL structure at 290 K. (B) The *ec*DHFR complex adopts the Met20 closed conformation and two rotamer states can be modeled for Met20 (both shown in stick representation), accompanied by unmodeled density. The bottom panel depicts how this electron density can be interpreted as a superposition of a “Gate Open” state that allows water into the active site and a “Gate Closed” state that occludes water. (C) The region composed of Met20, Pro21, and Trp22 adopts two conformations marked by distinct backbone conformations between Pro21 and Trp22 (blue and red arrows). (D) Kernel density estimates of the Trp22-*ϕ* dihedral from MD simulations in the context of a crystal lattice and a solvated water box, and dash marks to indicate the Trp22-*ϕ* dihedral in deposited structures of *ec*DHFR. The two states observed in (C) are shown with corresponding blue and red arrows, and the inset structure indicates the Trp22-*ϕ* dihedral. The 2*mFo − DFc* and *mFo − DFc* maps shown in (B) and (C) are carved within 1.5 Å and 3 Å, respectively, of the indicated residues for clarity.

Although ordered water is not observed in the active site of the Michaelis complex, the rotamer state of Met20 is hypothesized to regulate access of a water molecule to the N5 atom of DHF based on conformational heterogeneity in high-resolution structures [18]. Proton transfer directly from the solvent is further supported by molecular dynamics (MD) simulations and neutron diffraction [18–21]. Whereas proton transfer requires transient solvent access to the active site [18, 21], the presence of water near the N5 atom of DHF would destabilize the partial positive charge on the C6 carbon, inhibiting hydride transfer [22]. These observations, therefore, raise a more specific question: how does the enzyme regulate solvent access to tune the electrostatic environment of its active site to promote successive chemical steps with conflicting requirements—protonation which requires solvent access and hydride transfer for which solvent access is inhibitory.

Here, we apply new crystallographic methods to resolve conformational changes in *ec*DHFR, revealing rearrangements critical to the enzyme’s active site. First, by room-temperature X-ray diffraction, we observe extended conformational heterogeneity within the closed Met20 loop. By perturbing the active site with a modified substrate analog, we show direct coupling between the Met20 sidechain and the proton-donating water site. To assess the effect of larger-scale protein motions on the active site, we use new types of multi-temperature and electric-field stimulated [23] X-ray diffraction experiments. These methods resolve a surprising array of conformational motions—a global hinge motion that constricts the active site cleft and influences the Met20 sidechain, along with local networks of coupled backbone and sidechain motions affecting the active site. We validate this allosteric coupling by MD simulations and find that the protonated intermediate engages these motions by conformational selection to shield the active site from bulk solvent— a rapid rearrangement of the active site that follows substrate protonation to promote hydride transfer. We discuss several biological implications of this mechanism. For example, it explains a “dynamic knockout” mutant of *ec*DHFR—a mutant of which the effects on hydride transfer rate were proposed to result from altered dynamics alone, and not from a change in ground state structure [24]. We also describe how the mechanism appears to have constrained the evolution of the enzyme.

The approach taken here, combining advanced X-ray diffraction experiments with MD simulations, identifies global and local conformational dynamics that promote efficient catalysis. We expect that for many natural and designed proteins, this approach will similarly reveal important conformational rearrangements and answer fundamental questions about how these proteins work.

## Results

### The closed Met20 loop exhibits distinct substates

Structural, kinetic, and computational studies, combined with mutagenesis, have led to a basic understanding of how the active site of *ec*DHFR supports the chemical steps of catalysis. In this model, the Met20 sidechain regulates solvent access to the N5 atom of DHF to allow for substrate protonation (Fig. 1B) [17, 18]. To begin characterizing the conformational dynamics of the Michaelis complex, we used a widely employed model of the DHFR Michaelis complex with NADP^+^ and folate (FOL) as cofactor and substrate analogs, respectively, as the true Michaelis complex is not stable for the timescales necessary for crystallization [9]. The crystal form we used is also compatible with all steps of the catalytic cycle [9]. We first solved a structure of the model Michaelis complex to 1.04 Å at 290 K. Consistent with previous structures [9, 16, 18], the protein adopts the closed Met20 loop conformation, in which FOL and NADP^+^ are in close proximity (3.2 Å; Fig. 1B). Inspection of the electron density map (blue mesh, 2*mF_o_ − DF_c_*) near the Met20 sidechain shows electron density for two rotamers that differ in their *χ*_1_ dihedrals and the placement of the terminal methyl group. In addition, there is a large, 6.5*σ* peak in the difference electron density map between observed data and the refined model (green mesh, *mF_o_ −DF_c_*). This peak partially overlaps with one of the Met20 rotamer states (Fig. 1B), and can be identified as the proton-donating water by comparison with a previous X-ray diffraction study [18]. Together, these electron density features can be interpreted as a superposition of two Met20 sidechain conformations: a “gate open” Met20 rotamer can let water into the active site and a “gate closed” rotamer excludes water. This structure supports a solvent-gating role for Met20, and its analysis recapitulates the features observed by Wan *et al.* [18].

Our data, however, reveal additional conformational heterogeneity in the Met20 loop. The backbone amide between Pro21 and Trp22 adopts two distinct conformations, offset by approximately 90*^◦^* (arrows, Fig. 1C). These alternate backbone orientations can be thought of as substates of the closed loop conformation, and can be classified by the Trp22-*ϕ* dihedral angle with the two states centered at *−*150*^◦^* (blue arrow) and *−*75*^◦^* (red arrow). Although this heterogeneity has not been previously noted, we find a range of values for Trp22-*ϕ* consistent with these states in published structures of *ec*DHFR (Fig. 1D).

To assess whether the two substates represent dynamic exchange within the closed conformation of the Met20 loop, we ran MD simulations of the model Michaelis complex. When running the simulations in the context of the crystal lattice to recapitulate the impact of crystal contacts, we observe rapid sampling of transitions between the two substates, supporting that the crystallographic observation represents dynamic exchange. Based on classification using the Trp22-*ϕ* dihedral of each protein molecule in the simulation, we see that the substate at *−*75*^◦^* is populated approximately 2-fold more than the other substate (Fig. 1D). By fitting the simulation data to a Gaussian mixture model (see Methods) we can assign the populations as 66*±*3% and 34*±*3% (mean *±* standard error; N=72 trajectories), respectively, corresponding to ΔΔ*G ≈ −*0.4 kcal/mol (-0.7 *k_B_T*). This difference is similar to the relative density for the two states observed in the electron density map (Fig. 1C). This equilibrium also exists in MD simulations run in a waterbox, indicating that it is not an artifact of the crystal context. However, in a solvated system, the thermodynamics between the two substates inverts relative to that of the crystal lattice (Fig. 1D) with populations of 32 *±* 2% and 68*±*2% (mean *±* standard error; N=20 trajectories), respectively, corresponding to ΔΔ*G ≈* 0.4 kcal/mol (0.8 *k_B_T*).The crystal lattice therefore biases the thermodynamics between these states by about 0.8 kcal/mol (1.5 *k_B_T*).

### A modified substrate analog resolves the solvent gating mechanism

The model presented in Figure 1B suggests that the Met20 sidechain state regulates the occupancy of the proton-donating water. To test this hypothesis directly, we sought to bias the rotamer distribution of Met20 with a modified substrate analog, 10-methylfolate (MFOL). This compound has a methyl substituent on the N10 nitrogen (dashed circle in Fig. 2A) that makes close contact with the Met20 sidechain. We determined the structure of the *ec*DHFR:NADP^+^:MFOL complex to 1.14 Å (Table S1). As anticipated, this methyl group shifts the Met20-*χ*_1_ rotamer equilibrium (Fig. 2B). This structural change is accompanied by the appearance of an ordered water in the electron density map within 3.6 Å of the N5 nitrogen of MFOL (arrow in Fig. 2B), consistent with the location of the unmodeled difference density in Fig. 1C.

**Fig. 2.**
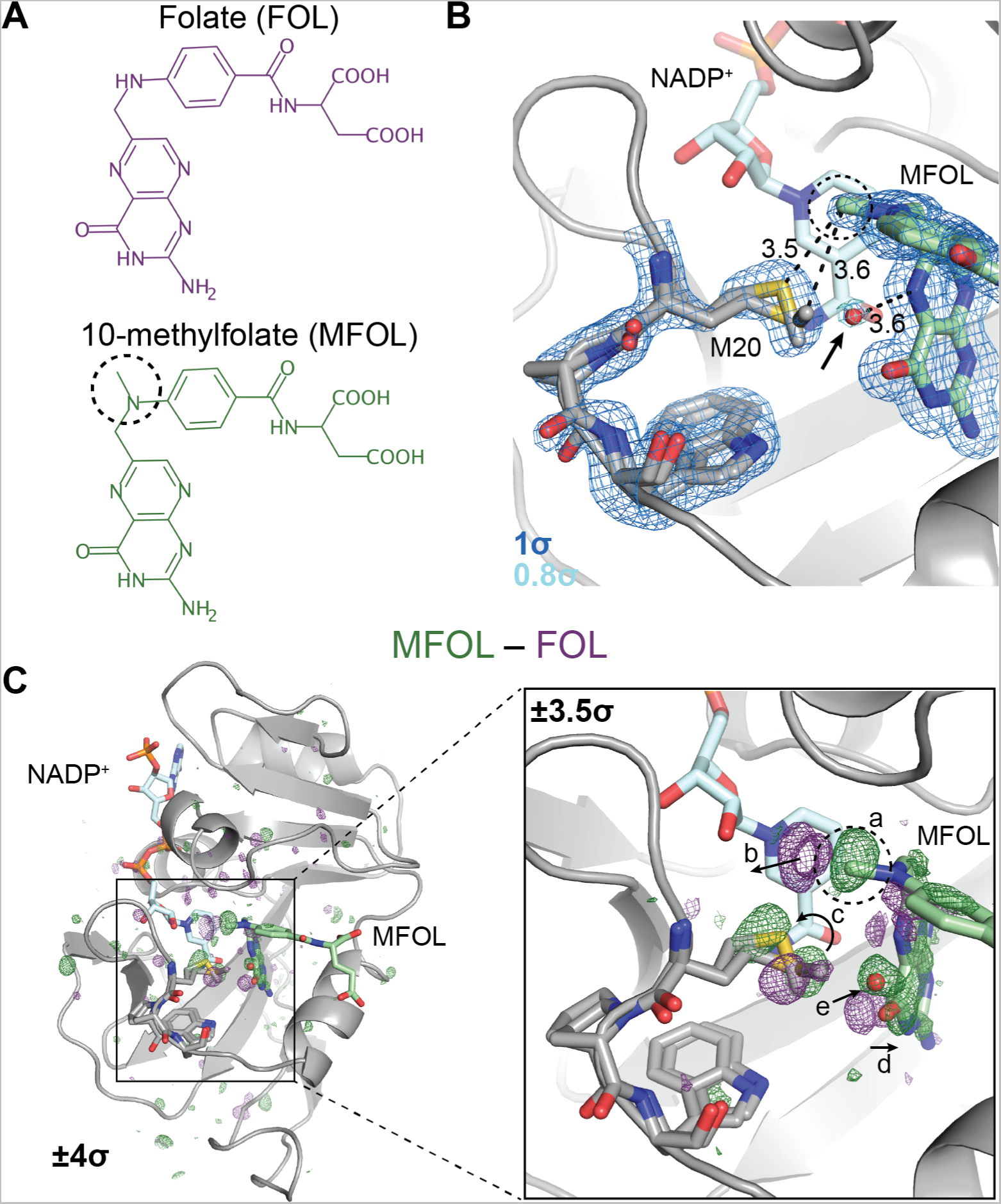
Ligand-dependent conformational changes illustrate Met20 solvent gating. (A) Chemical structures of folate (FOL) and 10-methylfolate (MFOL). (B) Refined structure and 2*mFo − DFc* electron density map of the *ec*DHFR:NADP^+^:MFOL complex. The 10-methyl group is in close contact with the Met20 sidechain, and a water (red sphere; indicated by an arrow) can be resolved within 3.6 Å of the N5 nitrogen of MFOL. The 2*mFo − DFc* map is contoured at 1*σ* (blue mesh; carved within 1.5 Å of shown atoms) and 0.8*σ* (light blue mesh; carved within 1.5 Å of shown water). (C) *F_MF OL_ −F_F OL_* isomorphous difference map, phased with the MFOL-bound model. The overview shows the difference electron density induced by the 10-methyl substituent (*±*4*σ*), and the inset highlights the structural differences observed in the active site (*±*3.5*σ*, carved within 3.0 Å of shown atoms). The added methyl group (label a) displaces an ordered water (label b), shifts the rotamer distribution of Met20 (label c), rotates the pterin ring (label d), and leads to the introduction of an ordered water near the N5 nitrogen (label e). The 10-methyl substituent is indicated with a dashed circle in each panel.

To identify the structural changes induced by the methyl substituent in more detail, we used the *F_MF OL_− F_F OL_* difference map, which can sensitively detect changes in electron density (Fig. 2C). Strong difference density is visible near the added methyl group (Fig 2C inset; labeled *a*). This 10-methyl group displaces two ordered waters from the folate-bound structure (labeled *b*), induces a shift in the Met20 rotamer distribution (labeled *c*), and causes the pterin ring to shift away from the Met20 residue (labeled *d*). Accompanying these changes, electron density for an ordered water increases near the N5 nitrogen (labeled *e*). That is, the 10-methyl substituent shifts the Met20 rotamer equilibrium, increasing solvent access to the N5 atom of the substrate.

### Multi-temperature diffraction resolves a global hinge motion

The structural changes observed in the 10-methylfolate complex validate the solvent-gating role of the Met20 sidechain, but were strongly localized near the 10-methyl substituent. Because the FOL-bound structure at 290 K and the MD simulations suggest additional conformational heterogeneity in the active site, we sought to bias the population of states of the enzyme using multi-temperature X-ray diffraction experiments. Because pre-existing equilibria that involve entropic change will be sensitive to temperature, these experiments can uncover correlated motions by observing structural states that change together as a function of temperature.

The earliest diffraction experiments to investigate the dependence of conformational heterogeneity on temperature used atomic displacement parameters as a reporter [25–27]. Since those early studies, multi-temperature X-ray crystallography has been applied to probe conformational changes caused by temperature with atomic detail in order to understand the dynamics of enzymes [28–30]. However, these experiments often probe a broad range of temperatures—from cryogenic to physiological—which can complicate analysis due to cryocooling artifacts and imperfect isomorphism [16]. Here, we collected 23 high-resolution datasets from crystals from 270 K to 310 K, in 10 K increments, including multiple datasets at each temperature to assess the uncertainty of any observations (Tables S2 to S6). We also inferred consensus datasets by combining data from the multiple crystals collected at each temperature (Fig. 3A and Table S7). To identify temperature-dependent structural changes within this physiological range, we adopted an automated refinement strategy yielding consistent models for each dataset. This approach enables detailed biophysical comparison across temperature.

**Figure 3:**
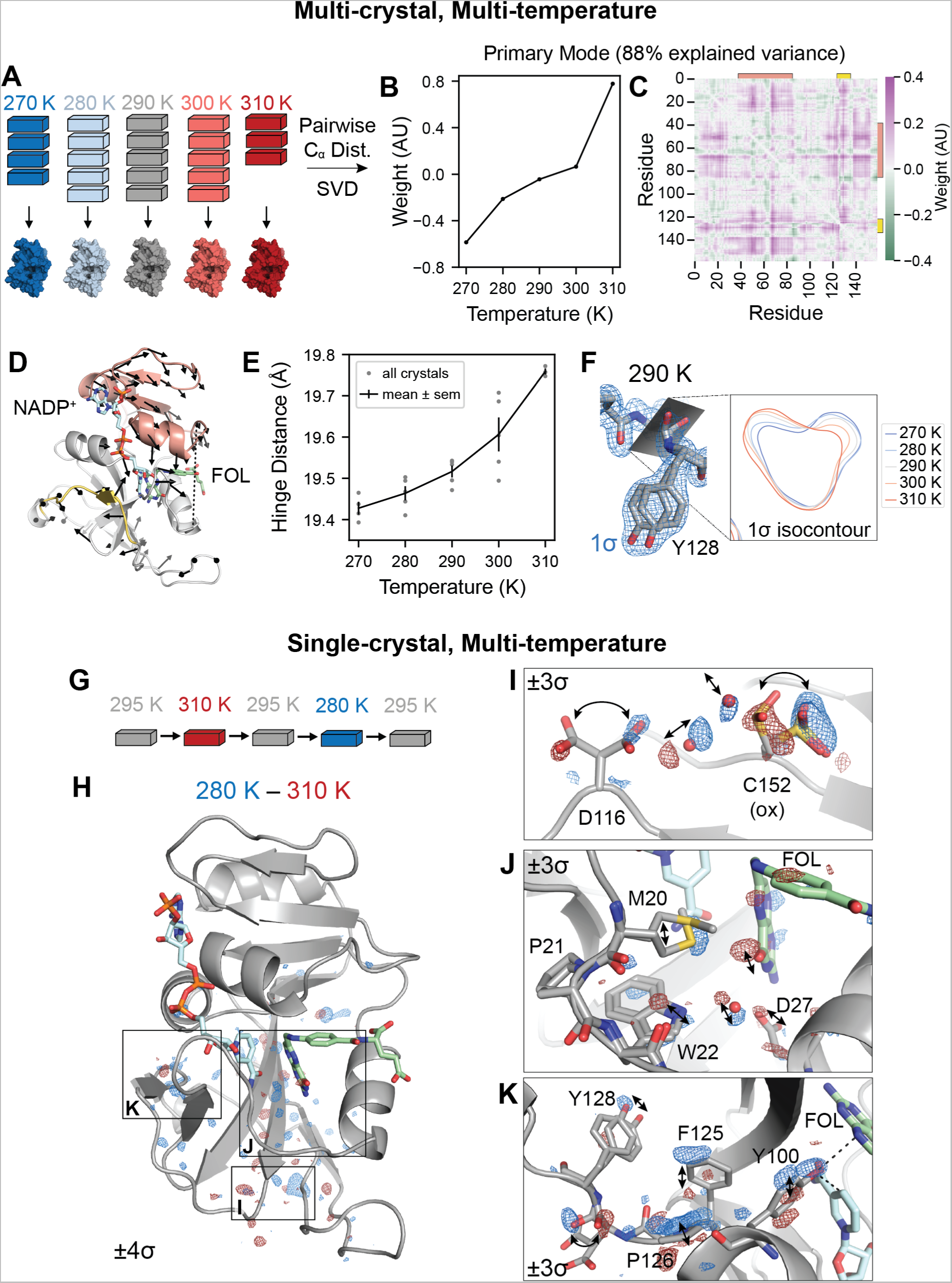
Multi-temperature experiments reveal a global hinge motion and local rearrangements. (A) Schematic of multi-crystal, multi-temperature diffraction experiment. (B) and (C) The primary structural mode from singular value decomposition (SVD) of the pairwise *C_α_* distances describes 88% of the variance among experimental structures. (B) Plot of the temperature dependence of the first left singular vector. (C) Heatmap of the contribution of each pairwise distance in the first right singular vector. Residues 38-88 are indicated with an orange bar and residues 120-130 are indicated with a yellow bar. (D) Structure of *ec*DHFR with arrows to depict displacements greater than 0.1 Å of *C_α_* atoms between 310 K and 270 K refined models. The arrows are enlarged 10-fold relative to the corresponding displacements. Residues 38–88 are shown in orange, residues 120-130 are shown in yellow, and the distance between Asn23-*C_α_* and Pro53-*C_α_* (hinge distance) is shown as a dashed line. (E) Plot of the hinge distance as a function of temperature. Data points are shown for each independent crystal and the mean *±* standard error at each temperature. (F) Structure and 2*mF_o_ − DF_c_* map for the 290 K consensus structure for Asp127 and Tyr128. The 1.0*σ* isocontour plot of the 2*mF_o_ − DF_c_* map in the plane of the backbone carbonyl is shown for the consensus structures at each temperature. (G) Schematic for the single-crystal, multi-temperature diffraction experiment. (H) Overview of the temperature-resolved isomorphous difference map between the 280 K and 310 K datasets. (I-K) Insets highlighting regions of the difference map. All maps are carved within 2 Å of the displayed atoms, and arrows highlight the structural changes. See also Figure S1.

To interpret overall conformational change, we computed the pairwise distances between the *C_α_* atoms in each refined structure for the consensus models at each temperature, and used singular value decomposition (SVD) to determine the primary temperature-dependent modes of structural change (see Methods for details). The resulting singular vectors describe the weights of the pairwise distances and temperature dependence for each structural mode. The first singular vector explains 88% of the variance of *C_α_* distances across datasets, and depends monotonically on temperature (Fig. 3B). The corresponding heatmap depicts the weight of each pairwise *C_α_* distance (Fig. 3C) and emphasizes two regions that correspond to residues 38-88 (orange bar) and residues 120-130 (yellow bar). These regions are colored on the structure of *ec*DHFR in Fig. 3D: residues 38-88, shown in orange, comprise the adenosine binding subdomain and residues 120-130, shown in yellow, span the end of the FG loop.

To illustrate the temperature-dependent motion corresponding to the first singular vector, Fig. 3D depicts the displacements in *C_α_* positions between the models refined to the 270 K and 310 K datasets. These are rendered as arrows for displacements greater than 0.1 Å and are enlarged 10x relative to the corresponding displacement. The arrows reveal a hinge motion that constricts the active site cleft. One of the strongest features in the pairwise distance heatmap corresponds to the distance between Asn23-*C_α_* and Pro53-*C_α_* (hereafter: hinge distance), which increases with temperature (Fig. 3E). Together, this analysis reveals a dominant, temperature-dependent global hinge motion that constricts the active site cleft by about 0.5 Å. Although this is a small-amplitude motion, the largest standard error in Fig. 3E is only 0.04 Å among replicate datasets.

In addition to the hinge motion, the region comprising residues 120-130 shows significant temperature dependence in Fig. 3C. In this region, Tyr128 adopts two shifted sidechain conformations, marked by distinct states for the amide backbone between Asp127 and Tyr128 (Fig. 3F). Accordingly, the refined electron density maps show a titration of density from one backbone configuration to the other as a function of increasing temperature, reaching equal occupancy at about 290 K (Fig. 3F).

### Temperature-resolved difference maps identify networks of correlated motions

The analysis of multi-crystal, multi-temperature diffraction experiments above identifies a global hinge-bending motion and shifts in the conformational equilibrium of the loop containing Tyr128. This approach works best to detect such graded shifts of the dominant conformation. Inspired by time-resolved diffraction experiments [31, 32], we sought to improve the detection of excited states by conducting single-crystal perturbation experiments, followed by analysis with isomorphous difference maps. In these experiments, we collected diffraction data at multiple temperatures from the same crystal (Fig. 3G). Difference maps obtained this way showed reproducible and remarkably sensitive results (see Methods and Figure S1).

The temperature-resolved difference maps obtained from single-crystal experiments reveal a range of conformational changes that were not readily detected by the multi-crystal, refinement-based analysis. The *F*_280*K*_ *−F*_310*K*_ isomorphous difference map is relatively flat in the adenosine binding subdomain, but exhibits regions of paired positive and negative difference density in the loop subdomain (Fig. 3H), which identify networks of temperature-dependent motion propagating through the enzyme, in addition to the large-scale hinge motion.

Three interesting regions of the protein have strong (*>* 5*σ*) peaks in the *F*_280*K*_ *− F*_310*K*_ difference map (Fig. 3H). As illustrated in Fig. 3I, the most significant difference map peak (10.3*σ*) involves the oxidized Cys152 sidechain and the nearby rotamers of Asp116. The paired difference density on the rotamers implies a correlated shift in their occupancy, which can be rationalized based on the corresponding movement of ordered water molecules found between these sidechains. A second network of temperature-dependent changes (5.6*σ* peak) runs through the active site including the Met20 loop (Fig. 3J). Paired difference density on the pterin ring of folate indicates that the ring settles deeper in the binding site with the constriction of the active site cleft. Asp27, which coordinates the pterin ring, shifts accordingly along with an ordered water bridging Asp27 and the Trp22 indole ring. Corresponding motions are observed in the Met20 loop itself, with a small shift in Trp22 and stronger density for the gate-open Met20 rotamer at lower temperature. Finally, the region from Phe125 to Tyr128 again shows significant temperature-dependent features in the difference map (5.5*σ*; Fig. 3K). The backbone amide between Asp127 and Tyr128 shows strong, paired difference density, consistent with the differences observed during refinement (Fig. 3F). The difference map, however, provides more detail, allowing the backbone carbonyl to be matched with the corresponding Tyr128 sidechain conformation based on their shared temperature dependence. Furthermore, strong difference density is observed for Pro126, Phe125, and Tyr100, highlighting an extended, contiguous network of temperature-dependent conformational changes that spans about 15 Å to the site of hydride transfer. Previous studies support the significance of these residues in catalysis. Tyr100 plays an important electrostatic role in hydride transfer [33], and the Y100F mutation decreases *k_hyd_* by ten-fold [17]. Similarly, double-mutant studies implicate Phe125 as part of a network of residues coupled to hydride transfer [34, 35]. In summary, single-crystal temperature-resolved diffraction experiments reveal detailed views of three extended networks of correlated motions that propagate throughout the enzyme and involve key active site residues.

### Electric-field-dependent constriction of the active site cleft

Although temperature can effectively bias conformational equilibria to observe correlated changes by X-ray diffraction, it impacts all states that differ entropically, possibly confounding a mechanistic interpretation of observed conformational changes. To further resolve the coupling between observed motions, we used electric-field-stimulated X-ray crystallography (EF-X). In an EF-X experiment, a strong electric field is used to apply force on the charges and local dipoles within a protein crystal to induce motions. These motions can then be observed by X-ray diffraction at room temperature (Fig. 4A). By using X-ray pulses at defined delays after the onset of the electric field, the induced dynamics can be followed with nanosecond temporal and sub-Ångstrom spatial resolution. EF-X has been used to study a PDZ domain, and the observed motions were consistent with proposed mechanisms of ligand-induced allostery [23]. Here, we used an updated apparatus for EF-X as shown in Figure 4B and S2A (see Methods for details). At each orientation of the crystal we collected 3 timepoints: an ‘Off’ reference timepoint in the absence of a high-voltage pulse, a 200 ns timepoint during a 3.5 kV pulse, and a 200 ns timepoint during a *−*3.5 kV pulse. To collect a complete dataset, we then rotated the sample, repeating the timepoints at each angle. This interleaved data collection ensures similar accumulated X-ray exposure for each dataset (Fig. 4C). The data collection statistics are presented in Table S11.

**Figure 4:**
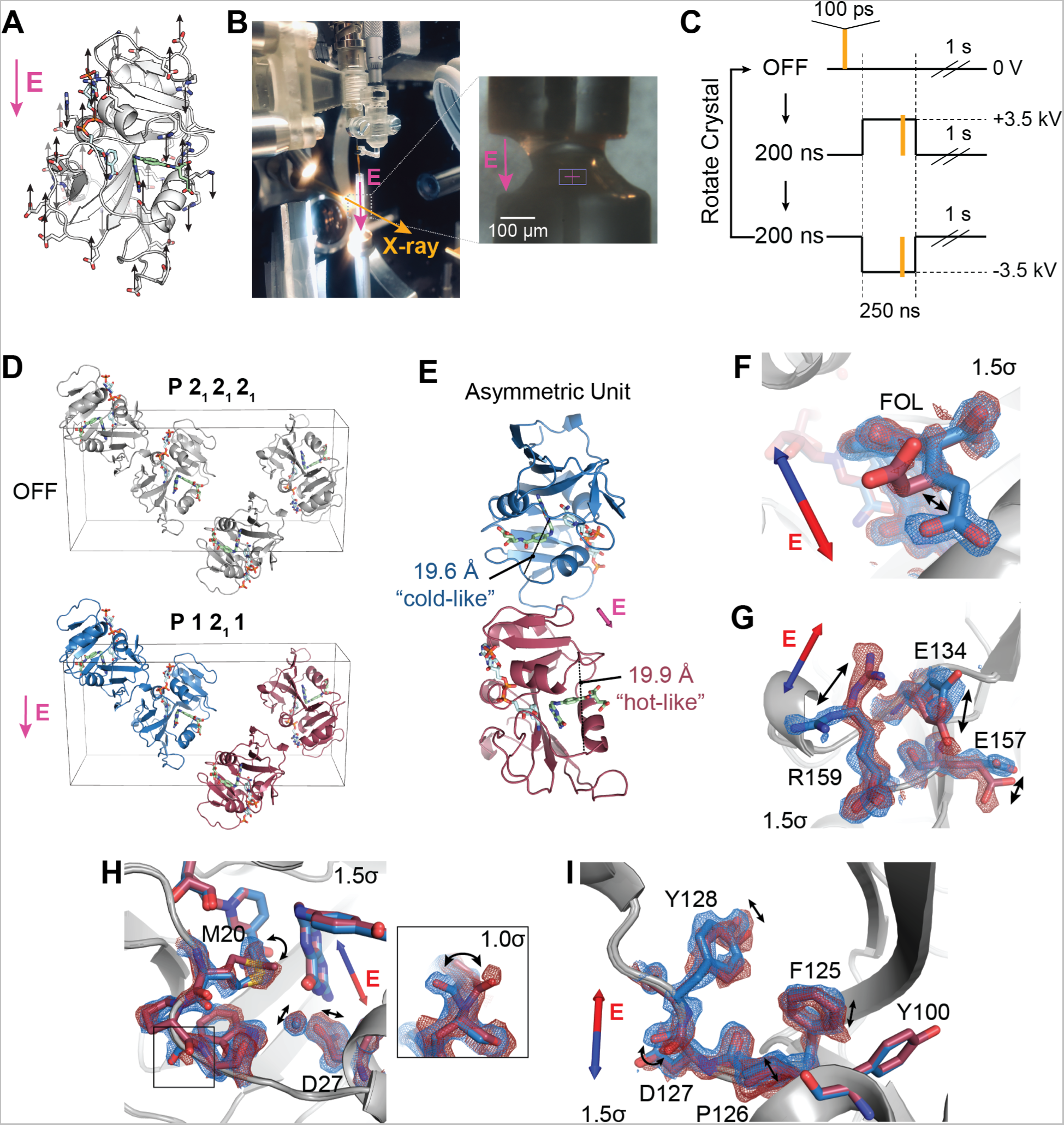
Electric-field-dependent structural changes recapitulate hinge motion and influence on active site residues. (A) Diagram of a possible pattern of force applied by an external electric field (E, in magenta) to *ec*DHFR based on the distribution of charged residues. (B) Photograph of the experimental apparatus for electric-field-stimulated X-ray crystallography (EF-X) at the BioCARS ID-B beamline; (inset) zoom-in showing an *ec*DHFR crystal between two electrodes. (C) Schematic of the data collection strategy, which included 3 consecutive X-ray pulses at each angle: OFF (no high voltage pulse), 200 ns into a +3.5 kV pulse, and 200 ns into a -3.5 kV pulse. The crystal was rotated after each sequence of 3 diffraction images in order to collect a complete dataset for each condition. (D) Unit cell of the *ec*DHFR crystal during the EF-X experiments. During the OFF images, the crystal is in the *P* 2_1_2_1_2_1_ spacegroup. The applied electric field along the *b*-axis alters the symmetry of the crystal, rendering the crystal in a *P* 12_1_1 spacegroup during the high voltage pulses, with two copies in the new asymmetric unit (ASU; copies shown in red and blue). (E) The ASU of the refined excited state model. The two copies in the ASU differ in hinge distance. The different copies of the protein are colored in red and blue as an analogy to the multi-temperature experiment; red represents the expanded active site cleft observed at hotter temperatures and blue represents the constricted cleft observed at colder temperatures. (F) to (I) Superposed models and 2*mF_o_ −DF_c_* maps from both protein molecules of the excited state ASU highlight electric-field dependent motion of charged groups. Blue and red arrows depict electric field vector for the blue and red models, respectively, and maps are contoured at 1.5*σ* and carved within 1.5 Å of shown atoms. (F) carboxylate sidechain of folate and (G) charged sidechains near the C-terminus demonstrate electric-field-dependent structural changes consistent with the formal charges of the residues. (H) Active site residues and Pro21 backbone carbonyl (inset; contoured at 1.0*σ*) differs between protein molecules. (I) Conformational changes among residues 125 to 128. Structural differences in panels F–I are also supported by composite omit maps, indicating that the results cannot be attributed to model bias (Fig. S3). See also Figure S2.

The high-voltage pulse applied in an EF-X experiment is directional. Copies of *ec*DHFR in the crystal’s unit cell are initially related by the symmetry operations of the *P* 2_1_ 2_1_ 2_1_ spacegroup. During the pulse, these copies experience the electric field, and therefore patterns of forces, in different orientations (Fig. 4D). In our case, two copies of *ec*DHFR experience the electric field in nearly the same direction (e.g., both blue copies) while the other two molecules (both red copies) experience the opposite field. The resulting deformations are therefore different for the red and blue copies. Notably, we can use the resulting symmetry breaking to confirm that there is significant signal in the experiment (see Methods for details; Fig. S2B).

To interpret the structural changes during the high-voltage pulse, we refined models of the induced excited states (see Methods for details). Significantly, the copies of the model Michaelis complex seeing the electric field in opposite direction refined to different hinge distances (19.6 Å for the ‘blue’ copy and 19.9 Å for the ‘red’ copy, Fig. 4E and S2C). These changes recapitulate the hinge motion observed using multi-temperature diffraction experiments (Fig. 3). Accordingly, we chose the color scheme for the two protein molecules to emphasize the comparison: the constricted copy is colored blue for “cold-like” and the extended copy is colored red for “hot-like”. The resulting electron density maps show clear electric-field dependent effects in which positively charged sidechains, like Arg159, move with the electric field, and negatively charged sidechains, like Glu134, move against the electric field (Fig. 4F, G), consistent with the expected movement of charge in an applied electric field. We also observe several shifts in the active sites of the two molecules, including motions of Asp27, the ordered water, and the sidechain rotamer of Met20 (Fig. 4H), as well as a flip in the backbone state of Pro21-Trp22 (Fig. 4H, inset). Because many residues in the Met20 loop lack a formal charge or significant charge dipole, these motions indicate conformational coupling of the Met20 loop with the rest of the enzyme. Furthermore, residues 125–128 display induced conformational rearrangements (Fig. 4I), similar to the conformational exchange observed in the multi-temperature experiment. Indeed, despite the very different perturbations being used, the sets of conformational changes observed in the active site and Tyr128 region for the multi-temperature and electric-field-dependent experiments are consistent in terms of the residues involved and the sign of the influence of the hinge distance. Together, this supports a common mechanism in which the global hinge motion is coupled to local rearrangements throughout the enzyme on the nanosecond timescale.

### Allosteric coupling of hinge motion to active site dynamics

MD simulations provide a means to directly validate the mechanistic model that the hinge motion allosterically regulates the local conformational equilibria in the active site. Specifically, we can bias the hinge distance in simulation using an imposed distance restraint to observe its impact on other observables in the protein. To do so, we applied a distance restraint across the active site cleft with equilibrium values chosen to span the crystallographically observed range (Fig. 5A). We ran 100 independent, 100 ns MD simulations at each hinge distance. These restraints successfully biased the sampled conformations to particular widths of the active site cleft (Fig. 5B). In response, the population of states of the Met20 loop backbone changes monotonically (Fig. 5C, using the Trp22-*ϕ* backbone dihedral as a reporter). Similarly, with increasing hinge distance the Met20 sidechain shifts its rotamer distribution, as reported by a decrease of the population of the *χ*_1_ dihedral around *χ*_1_ = *−*160*^◦^* (Fig. 5D). This change is consistent with the multi-temperature experiment, in which the Met20-*χ*_1_ of approximately *−*160*^◦^* was more populated at lower temperature (shorter hinge distance; Fig. 3J). This is also consistent with the EF-X results, in which the copy with a shorter hinge distance favored the Trp22 backbone and Met20 rotamer states observed in MD (Fig. 4H). These simulation results, therefore, corroborate the crystallographic analysis and confirm that the width of the active site cleft is allosterically coupled to the occupancy of the Met20 loop substates.

**Fig. 5.**
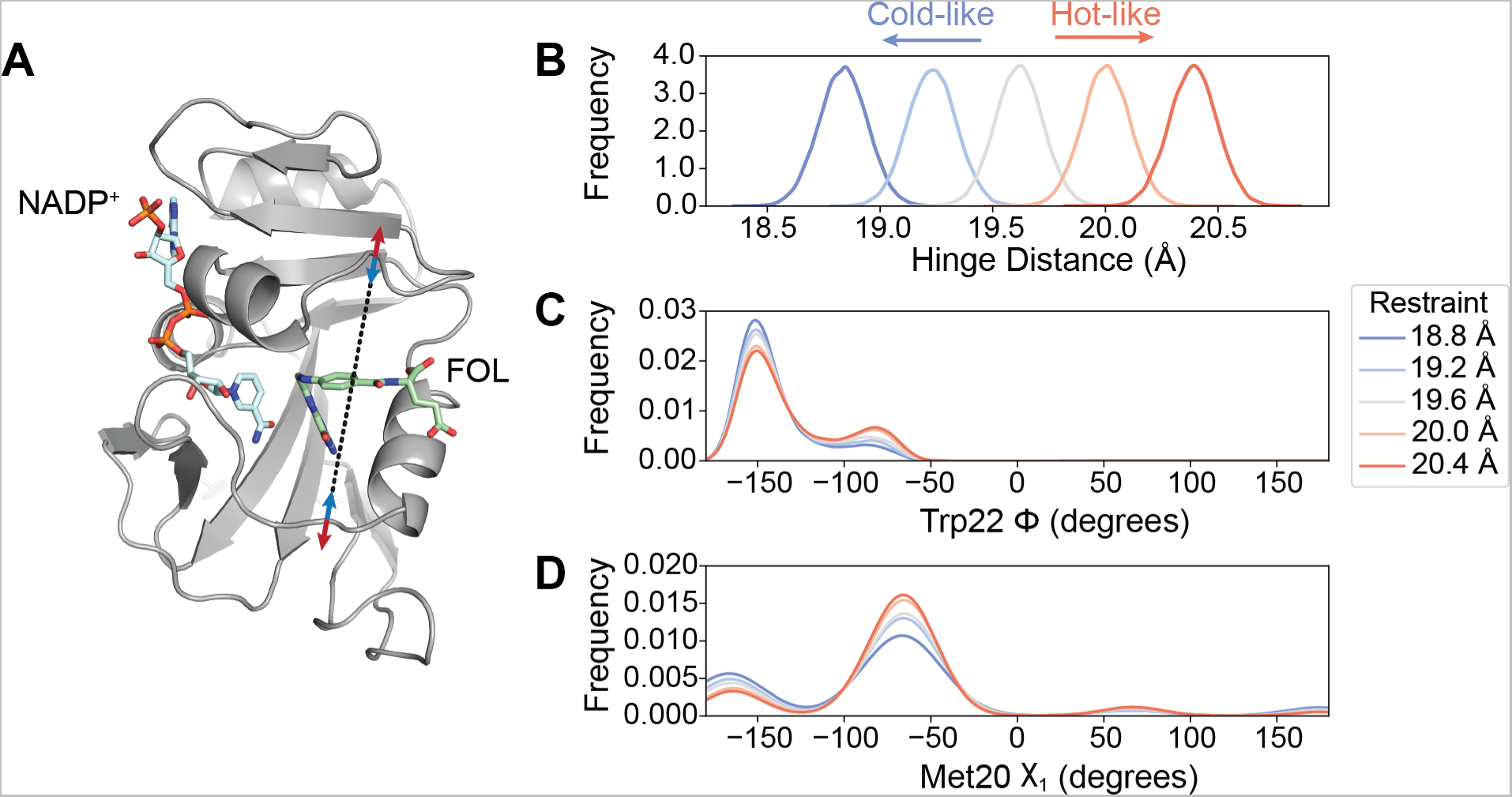
MD simulations validate the influence of hinge motion on the substates of the closed Met20 loop. (A) Simulation model of *ec*DHFR:NADP^+^:FOL highlighting the distance restraint applied in MD simulations between Asn23-*Cα* and Pro53-*Cα* (black dashed line) to model the effects of constricting (blue arrows, cold-like by analogy to the multi-temperature experiment) or expanding (red arrows, hot-like) the active site cleft. Kernel density estimates of the (B) hinge distance being restrained, (C) the Trp22-*ϕ*, and the (D) Met20-*χ*_1_ dihedrals. The Trp22-*ϕ* and Met20-*χ*_1_ dihedrals, which report on the Met20 closed substates, show a monotonic response to the distance restraint. The kernel density estimates were produced from 100 independent simulations of 100 ns duration at each restraint distance.

### Substrate protonation regulates active site solvent access

Do these global and local active site motions impact catalysis? As described, reduction of dihydrofolate involves two sequential steps: substrate protonation and hydride transfer. To address the effect of protonation on the reactive Michaelis complex (DHFR:NADPH:DHF), we ran MD simulations of the deprotonated and N5-protonated complexes. Statistical distributions of key structural parameters are shown in Figure 6. Upon protonation, the average hinge distance decreases by approximately 0.5 Å and the Trp22-*ϕ* equilibrium is further shifted towards the state near *−*150*^◦^*. This combination of changes recapitulates the allosteric mechanism identified above, and indicates that substrate protonation engages this dynamic mode.

**Fig. 6.**
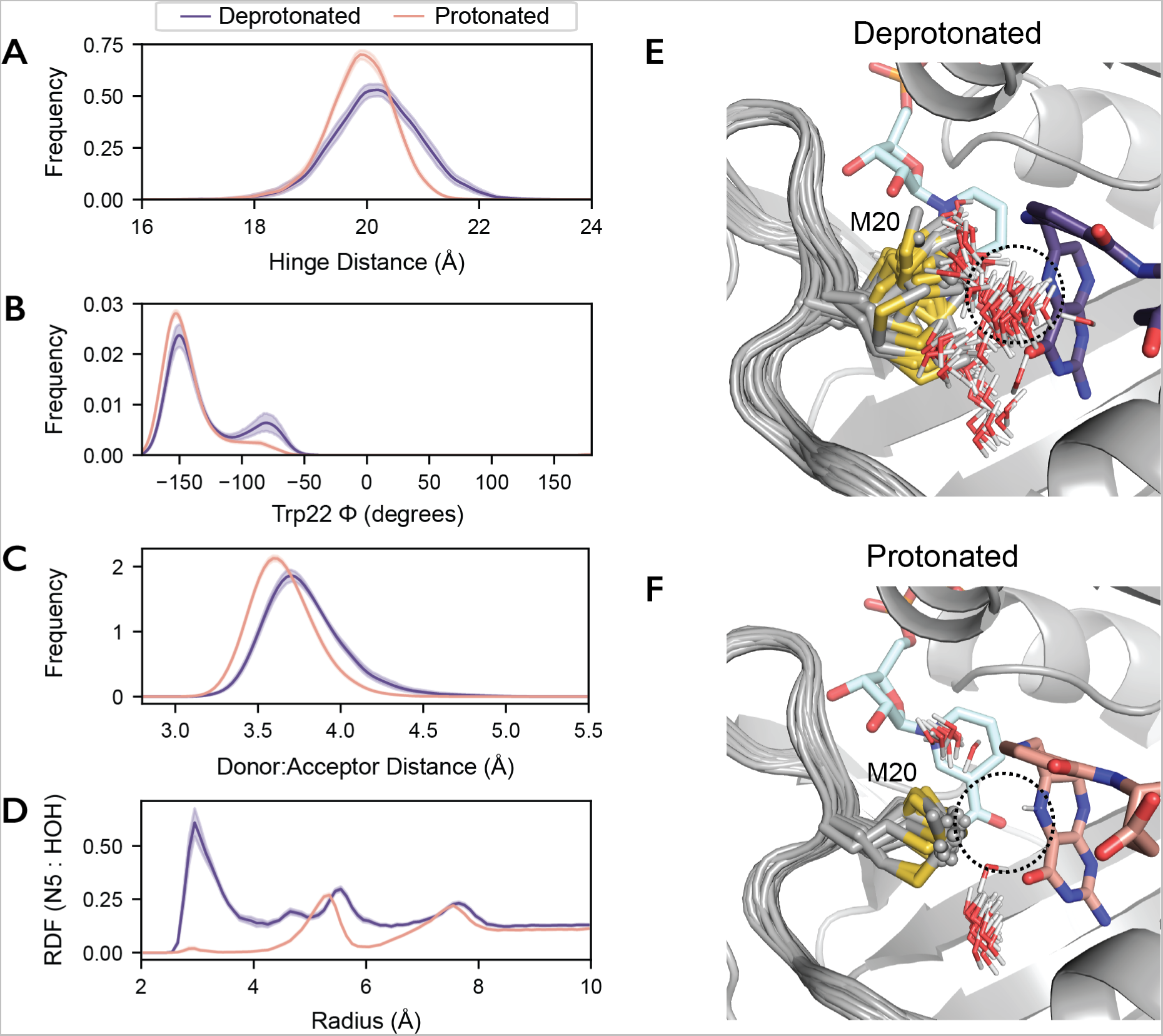
Protonation of the substrate orders the Met20 sidechain in the Michaelis complex. 50 independent MD simulations of the *ec*DHFR:NADPH:DHF complex, with and without protonation of the N5 nitrogen, were run for 100 ns each. Kernel density estimates of the (A) hinge distance, (B) Trp22-*ϕ*, (C) donor-acceptor distance for hydride transfer change upon protonation of the substrate. These kernel density estimates were computed for each trajectory independently and the mean and 95% confidence interval is shown for each condition. (D) The density of water around the N5 nitrogen of DHF as a function of distance from the N5 nitrogen (radial distribution function; RDF) mean and 95% confidence interval are shown. The first 50 frames (20 ns) from one trajectory are superimposed for the (E) deprotonated and (F) protonated substrate, depicting the Met20 sidechain and all waters within 4.5 Å of the N5 nitrogen of DHF. Only the initial frame is depicted for DHF and NADPH for visual clarity.

**Fig. 7.**
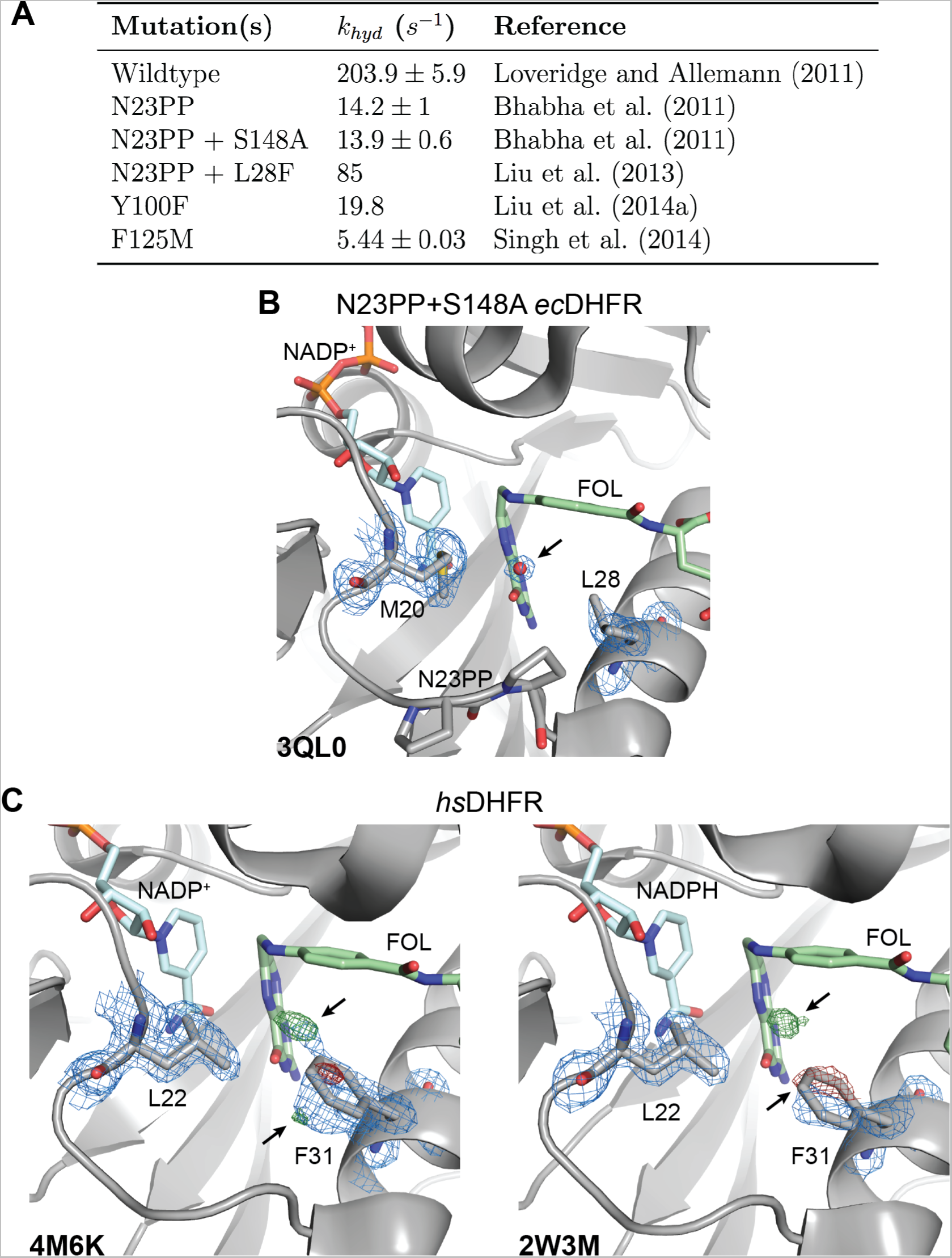
Functional importance and conservation of solvent gating in DHFR. (A) The rate of hydride transfer, *k_hyd_*, for selected mutants of *ec*DHFR. (B) The structure of the N23PP/S148A mutant of *ec*DHFR (PDB: 3QL0) shows wellsupported density for an ordered water in the 2*mFo − DFc* map (blue mesh; 1.5*σ*). (C) Structures of human DHFR (PDB: 4M6K and PDB: 2W3M, molecule B) have unmodeled density consistent with partial-occupancy water within 3.5 Å of the N5 nitrogen of FOL and evidence of an alternate rotamer for Phe31 (*mFo − DFc*; green/red mesh; *±*3.5*σ*). A single rotamer is supported for Leu22 in the 2*mFo − DFc* maps (blue mesh; 1.0*σ*) suggesting that Phe31 instead serves as the solvent-gating residue in the human enzyme. Although only molecule B is presented for the 2W3M deposited structure, similar features are observed in both protein molecules of the asymmetric unit. See also Figure S4.

The donor-acceptor distance for hydride transfer also decreases upon protonation (Fig. 6C). This distance is the primary determinant of hydride transfer [36], and the change is consistent with the increase in the partial charge assigned to the C6 of DHF upon protonation. Interestingly, protonation of the N5 nitrogen also effectively eliminates water from its proximity by ordering the Met20 sidechain. Indeed, the radial distribution function of water molecules around the N5 nitrogen indicates very little occupancy of the proton-donating water site after protonation (Fig. 6D), consistent with findings in complementary simulation-based studies [37, 38]. To visualize this change in the organization of the active site, we superpose frames from the trajectories. Overlaying 20 ns of one representative trajectory shows heterogeneity in the Met20 rotamer and frequent occupancy of the water site (dashed circle) for the deprotonated substrate (Fig. 6E), whereas the protonated substrate coordinates the Met20 rotamer that occludes the water site (Fig. 6F).

Experimentally, we also observed that the network involving Tyr128, Phe125, and Tyr100 exhibits pronounced temperature dependence (Fig. 3C) and motions extending from Tyr128 to the active site residue Tyr100 (Fig. 3K). This network did not respond to variation in hinge distance in MD simulations (Fig. S4A) but does respond to substrate protonation in MD (Fig. S4B). Most likely, then, this network of residues contributes to electrostatic remodeling of the active site in response to protonation independently from the enzyme’s hinge motion.

In summary, before protonation the active site has a pre-existing equilibrium of states that permits solvent access to the N5 nitrogen. This equilibrium is allosterically coupled to the width of the active site cleft. This dynamic architecture allows the enzyme to quickly reorganize the active site in response to protonation of its substrate. This rearrangement facilitates hydride transfer by polarizing the C6 carbon, shortening the donor-acceptor distance, and inhibiting the competing deprotonation reaction by excluding bulk solvent, consistent with a proposal by McTigue *et al* [22].

## Discussion

By a combination of new X-ray diffraction experiments and analysis, we resolved the correlated motions of an enzyme in atomic detail. Using room-temperature diffraction, we first identified extended conformational heterogeneity in the enzyme’s active site loop (Fig. 1C). We then used a substrate mimetic to demonstrate that the Met20 sidechain directly regulates solvent access to the active site (Fig. 2C). Multi-temperature and EF-X experiments then uncovered a global hinge motion that constricts the enzyme’s active site and local networks of conformational rearrangements throughout the enzyme (Fig. 3 and 4). MD simulations confirmed that the hinge motion has a direct allosteric effect on conformational equilibria within the active site (Fig. 5). This coupling enables the protonated substrate to rapidly select an active site arrangement that favors the subsequent hydride transfer step over deprotonation (Fig. 6). The result is a model of catalysis by *ec*DHFR in which the product of the first chemical step (a reaction intermediate) drives rapid rearrangments in the active site by conformational selection to favor the second chemical step. That is, the enzyme is wired to undergo conformational change in response to completion of the first chemical step, just like it does after substrate binding, product formation, and product release, a view that naturally extends the notion of a dynamic free energy landscape as the organizing principle of enzyme catalysis [11].

### Functional significance of solvent gating in *ec*DHFR

Our work validates the proposed solvent-gating role of Met20 and resolves conformational dynamics in *ec*DHFR that allosterically regulate the organization of the active site in response to substrate protonation. But, how important is proper solvent gating for hydride transfer? An important case study for the role of the Met20 loop in catalysis is the N23PP *ec*DHFR mutant (and the related N23PP/S148A mutant) that introduces the double proline insertion found in the human enzyme. This mutation decreases the rate of hydride transfer (*k_hyd_*) by approximately 15-fold (Fig. 7A) with little apparent change in the overall structure. Because relaxation-dispersion experiments showed that this variant no longer displays millisecond-dynamics of the Met20 loop, Bhabha *et al.* concluded that these motions influence the chemical step(s) of catalysis in *ec*DHFR and classified the mutant as a “dynamic knockout” [24]. Adamczyk *et al.* disputed this conclusion with arguments about the importance of electrostatic preorganization and MD simulations showing no productive relationship between a putative coordinate for millisecond dynamics of the Met20 loop and the energy barrier for hydride transfer [39]. Loveridge *et al.* found that although the insertion mutant showed a reduced rate of hydride transfer, the corresponding kinetic isotope effect and its temperature dependence were largely unaffected [36]. They interpreted this as evidence that the mutation does not alter direct dynamic contributions to hydride transfer. Based on our work, we believe that the N23PP mutation impedes the solvent-gating activity of *ec*DHFR: Close inspection of the active site in the published N23PP/S148A *ec*DHFR structure [24] shows that a water occupies the site typically occluded by Met20 in the wildtype enzyme (Fig. 7B; PDB: 3QL0). The proline insertion increases the spacing between Met20 and the subsequent *α*-helix by about 0.3 Å (measured from Met20-*C_α_* to Leu28-*C_α_*), such that the methionine sidechain no longer blocks solvent access to the substrate, trapping the protein in a gate-open state that is less competent for hydride transfer. In this view, the solvent-gating function of Met20 mechanistically underpins the effect of the N23PP mutation, providing a structural explanation for the prior notion that the insertion disrupts the electrostatic environment of the *ec*DHFR active site [39].

Consistent with this inference, about six-fold of the catalytic activity of N23PP can be rescued by the point mutation L28F (Fig. 7A), which introduces a larger residue on the adjacent *α*-helix [40] and likely partially restores the capability to shield the substrate from solvent. These results are consistent with a central role for solvent-gating in enhancing the hydride transfer rate of *ec*DHFR.

### Dynamic modes and solvent-gating function are conserved in DHFR

Evolutionary conservation provides further perspective on the importance of the observed motions as DHFR homologs catalyze the same reaction and face similar challenges. The hinge motion characterized here within the model Michaelis complex resembles the conformational changes observed between substrate and product ternary complexes [41] in terms of its associated changes in pairwise-distance between *C_α_* atoms (Fig. 3C). The latter motion reflects a small (*<*1 Å) hinge motion, and has been described as a subdomain rotation that alters the width of the active site cleft [9]. Notably, the human homolog exhibits a substantially larger hinge motion (*∼*3 Å) upon product release [41], which was postulated to facilitate cofactor exchange in versions of DHFR with a more rigid Met20 loop [41]. Consistently, the occluded state of the Met20 loop, which facilitates cofactor release in *ec*DHFR, has not been observed in eukaryotic DHFRs [9, 10, 41].

Indeed, the Met20 loop of human DHFR does not exhibit the conformational flexibility observed for the *E. coli* enzyme [9], and the analogous residue to Met20, Leu22, has well-resolved density for a single conformation in models of the human DHFR Michaelis complex (Fig. 7C). Strikingly, however, the differences between the modeled and observed electron density (*mF_o_ − DF_c_*) for two previously deposited structures of human DHFR both show clear evidence of an excited state rotamer of the Phe31 sidechain (Fig. 7C). Accordingly, there is unmodeled positive difference density near the N5 nitrogen of folate, suggesting partial occupancy by a proton-donating water as observed for *ec*DHFR (Fig. 1B). Together, these observations strongly suggest that in human DHFR Phe31 is functionally analogous to *ec*DHFR Met20, rather than the structurally homologous Leu22 residue. This functional analogy was first proposed by McTigue *et al.* [22] and implies that solvent-gating is functionally conserved in the active sites of DHFR enzymes. Considering these structural observations along with partial functional rescue of the N23PP mutation by the L28F mutation in the *E. coli* enzyme, this suggests a mechanistic basis for the appearance of both mutations at a similar point in evolutionary history [40].

### Identifying functional networks of residues

The temperature-resolved difference maps produced in this work allowed us to trace networks of correlated structural changes across multiple residues. We observe the pterin ring of folate adopt a deeper binding pose in the active site with corresponding movement of the critical active site residues Met20, Trp22, Asp27, and an ordered water molecule (Fig. 3J). EF-X corroborates this network of correlated rearrangements (Fig. 4I), which can mechanistically explain the allosteric coupling we observe between the hinge motion and the active site (Fig. 5). Similarly, a network of functionally important residues including Tyr128, Phe125, and Tyr100 moves in response to perturbations (Fig. 3K; Fig. 7A). Difference map-based analysis of these diffraction experiments therefore now provides the sensitive detection of correlated motion needed to develop mechanistic models. The atomic and temporal resolution of these experiments naturally complement MD simulations in the development and testing of structural hypotheses.

In summary, the work presented here used ligand-, temperature-, and electric-field-dependent X-ray diffraction experiments and MD simulations to resolve a conserved dynamic mode that allosterically influences local conformational equilibria in the active site of *E. coli* DHFR. This reveals an enzyme with dynamics primed to respond to the protonation of its substrate. We believe the approach presented here will have broad application. The protein crystals we used are equivalent to those used for decades (for example in refs. [9, 15, 16]). However, the advances described here, building on improvements in hardware [23, 42], data collection strategies [7, 43], and analysis methods [44–46], enabled elucidation of the correlated motions of an enzyme in atomic detail. We expect the presented methods and strategy will likewise permit identification of the motions that underlie the function of a wide range of proteins, promoting the development of new mechanistic models to explain protein function and its allosteric regulation.

## Acknowledgements

We thank Drs. B. Correia, S. Eddy, R. Gaudet, and R. Losick for comments on the manuscript. D.R.H. is supported by the Searle Scholarship Program (SSP-2018-3240), a fellowship from the George W. Merck Fund of the New York Community Trust (338034), and the NIH Director’s New Innovator Award (DP2-GM141000). J.B.G. was supported by the National Science Foundation Graduate Research Fellowship under Grant No. DGE1745303. K.M.D. holds a Career Award at the Scientific Interface from the Burroughs Wellcome Fund. M.A.K. is supported by the NSF-Simons Center for Mathematical and Statistical Analysis of Biology at Harvard (award number #1764269) and the Harvard Quantitative Biology Initiative. Use of the Stanford Synchrotron Radiation Lightsource, SLAC National Accelerator Laboratory, is supported by the U.S. Department of Energy, Office of Science, Office of Basic Energy Sciences under Contract No. DE-AC02-76SF00515. The SSRL Structural Molecular Biology Program is supported by the DOE Office of Biological and Environmental Research, and by the National Institutes of Health, National Institute of General Medical Sciences (including P41GM103393). In addition, this research used resources of the Advanced Photon Source, a U.S. Department of Energy (DOE) Office of Science User Facility operated for the DOE Office of Science by Argonne National Laboratory under Contract No. DE-AC02-06CH11357. Use of BioCARS was also supported by the National Institute of General Medical Sciences of the National Institutes of Health under grant number P41 GM118217. The time-resolved set-up at Sector 14 was funded in part through a collaboration with Philip Anfinrud (NIH/NIDDK). The contents of this publication are solely the responsibility of the authors and do not necessarily represent the official views of NIGMS or NIH.

## Author Contributions

J.B.G., K.M.D., and D.R.H. conceived the project. C.J.S. and D.E.B. purified the protein, and J.B.G., K.M.D., and D.E.B. crystallized the *ec*DHFR complexes. J.B.G., K.M.D., and D.R.H. planned the multi-temperature diffraction experiments, and J.B.G., D.E.B., M.A.K., and S.R. conducted the data collection. J.B.G., K.M.D., D.E.B., M.A.K., I.K., R.W.H., and D.R.H. developed and conducted the EF-X experiments. J.B.G. analyzed the diffraction data for all experiments. J.B.G. ran and analyzed the molecular dynamics simulations. The manuscript was written with feedback from all authors.

## Data and Code Availability

All structures determined in this study have been deposited in the Protein Databank (PDB) with IDs: 8DAI, 5SSS, 5SST, 5SSU, 5SSV, 5SSW, 7FPL, 7FPM, 7FPN, 7FPO, 7FPP, 7FPQ, 7FPR, 7FPS, 7FPT, 7FPU, 7FPV, 7FPW, 7FPX, 7FPY, 7FPZ, 7FQ0, 7FQ1, 7FQ2, 7FQ3, 7FQ4, 7FQ5, 7FQ6, 7FQ7, 7FQ8, 7FQ9, 7FQA, 7FQB, 7FQC, 7FQD, 7FQE, 7FQF, 7FQG, 8G4Z, and 8G50, as referenced in the Supplementary Tables. Python and PyMOL scripts for generating figures, along with (difference) electron density maps are deposited in Zenodo (https://doi.org/10.5281/zenodo.7634123). Crystallographic analyses make use of `reciprocalspaceship` and `rs-booster`, which are available from https://rs-station.github.io/. The force-fields, starting models, and scripts for reproducing the molecular dynamics trajectories are included in the Zenodo deposition.

## Methods

### Protein Purification and Crystallization

We expressed, purified, and crystallized *ec*DHFR as described previously [43], with one modification. In order to purify *ec*DHFR for the complex with 10-methylfolate, we modified the methotrexate-affinity chromatography to include a wash with 200 mM potassium phosphate buffer (pH 6.0) with 1 M potassium chloride, 1 mM ethylenediaminetetraacetic acid (EDTA), and 1 mM dithiothreitol (DTT) and elution the protein using a linear gradient with 50 mM potassium borate buffer (pH 10.15) and 2 M potassium chloride. The high pH, high salt elution was necessary to avoid contamination of the purified protein with bound folate. We used crystals of the model of the Michaelis complex, *ec*DHFR:FOL:NADP^+^, for the multi-temperature X-ray diffraction experiments and the electric-field-stimulated X-ray diffraction (EF-X) experiments. We co-crystallized the 10-methylfolate (No. 16.211, Schircks Laboratories) complex using the same conditions as the *ec*DHFR:FOL:NADP^+^ complex [43].

### Monochromatic Data Collection

We collected the 10-methylfolate complex and multi-temperature datasets presented in this work at the Stanford Synchrotron Radiation Lightsource (SSRL) beamline 12-1 at the SLAC National Accelerator Laboratory. We collected the data during three beamtime allocations on July 20, 2021; November 10, 2021; and May 7, 2022. We looped all crystals at Harvard University using the MicroRT system (MiTeGen) for room-temperature data collection, and shipped the looped crystals to SSRL 12-1 using the SSRL *in situ* Crystallization Plate (M-CP-111-095, Crystal Positioning Systems) and a thermal shipping container to maintain the samples at 277 K. The specialized plate was used for compatibility with the robotic sample handling at SSRL 12-1, which supported remote data collection at regulated temperatures and high humidity [47]; see also https://www-ssrl.slac.stanford.edu/smb-mc/content/users/manuals/remote-access-at-elevated-temperatures-and-controlled-humidity.

For all monochromatic diffraction experiments we used helical data acquisition, translating along the long-axis of the rod-shaped crystals to best distribute the radiation dose among the crystal volume. Unless otherwise noted, the beam size was set to 50 *×* 50 µm^2^ and 0.2% transmission. On average, the crystals were 75 *×* 75 *×* 500 µm, and we collected 1440 images with a 1*^◦^* oscillation angle and a 0.2 s exposure time. SSRL BL12-1 is equipped with an Eiger 16M detector (Dectris) with a pixel size of 75 µm^2^. We began each crystal at 295 K, and adjusted the environmental temperature to the desired set point at a ramp rate of approximately 2 *^◦^*/min.

### 10-methylfolate Complex

We collected the diffraction data for the 10-methylfolate complex with a beam size of 50 *×* 7 µm^2^ at 13.00 keV, a detector distance of 160 mm, and at 285 K.

### Multi-temperature Diffraction Experiments

To investigate the conformational changes in DHFR across a range of physiological temperatures, we collected 4 datasets at 270 K, 5 datasets at 280 K, 5 datasets at 290 K, 1 dataset at 295 K, 5 datasets at 300 K, and 3 datasets at 310 K. For these experiments, we collected a single dataset at the desired temperature from each crystal, using an incident beam energy of 15.00 keV and a detector distance of 160 mm. To facilitate the use of isomorphous difference maps to identify structural differences, we also collected multiple datasets at different temperatures from the same crystal. For one crystal, we collected successive datasets at 295 K, 280 K, 295 K, 310 K, and 295 K, and for another crystal we collected the reversed series at 295 K, 310 K, 295 K, 280 K, and 295 K. The repeated measurements at 295 K allowed us to assess hysteresis and to rule out radiation damage, as indicated by the relatively flat isomorphous difference maps from successive datasets.

### Data Reduction, Scaling, and Structure Refinement

We used *DIALS* to find and index strong spots, refine the experimental geometry, and integrate each dataset at each temperature [44]. Each dataset was processed independently, using default parameters in *DIALS*. During indexing we provided the space group, *P* 2_1_2_1_2_1_, and used local index assignment (index.assignment.method=local). This improved the indexing rate by reducing the sensitivity to small crystal motions during the course of helical data acquisition. Following geometry refinement, the residuals for spot prediction were approximately 0.2-0.3 px (RMSD).

The relative scale of each dataset is an important consideration when using difference maps to visualize conformational changes between conditions. We used dials.scale with a common reference dataset, collected during the same day at 295 K, to ensure a consistent relative scale across all of our data [48]. In addition to scaling and merging each dataset individually, we scaled and merged data collected at the same temperature from multiple crystals to refine single, representative structures for each temperature. High-resolution cutoffs were always chosen such that the half-dataset correlation coefficient of the highest resolution bin was greater than 0.3 [49]. In all cases, the high resolution cutoff was *<* 1.35 Å, and the majority of the crystals diffracted to between 1.05 and 1.15 Å.

Due to the large number of diffraction datasets involved in this study, we chose an automated structure refinement protocol. We used `phenix.refine` [50] to refine occupancies, anisotropic B factors for all non-hydrogen atoms, and reciprocal space XYZ refinement to improve the atomic coordinates. Ligand geometry restraints for NADP^+^, folate, 10-methylfolate, and oxidized cysteine (cysteine sulfinic acid) were generated using `phenix.elbow` using default parameters. Due to the high degree of similarity between each dataset, we initialized each refinement run by isomorphous replacement, and we found ten macrocycles to be sufficient to converge the refinement R factors. Importantly, to ensure that R-factors were comparable between runs, we used a common R-free set composed of 5% of the unique reflections.

### Analysis of Multi-crystal, Multi-temperature Experiment

To identify temperature-dependent structural changes from refinement, we analyzed changes in pairwise distances between refined C*_α_* coordinates. For residues refined with alternate conformations, only the highest occupancy conformer was included in the analysis. We used the *SciPy* library [51] to compute the pairwise distances between coordinates. These distances were treated as features and computed for the consensus structures at each temperature, yielding a *N×d* matrix with *N* datasets and *d* features. To prioritize analysis of how the structures differed, we subtracted the mean of each pairwise distance from the corresponding rows of the matrix. We then used singular value decomposition in *NumPy* [52] to analyze the primary temperature-dependent mode among the datasets.

### Isomorphous Difference Maps

This work presents weighted isomorphous difference maps across temperatures and between different ligand-bound complexes. These maps used difference structure factor amplitudes, *|*Δ*F_H_ |*, given by

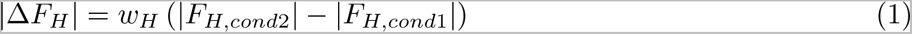

where *|F_H,cond_*_1_*|* and *|F_H,cond_*_2_*|* are the merged structure factor amplitudes for the first condition and second condition, respectively, and *w_H_* are weights defined as follows [53]:

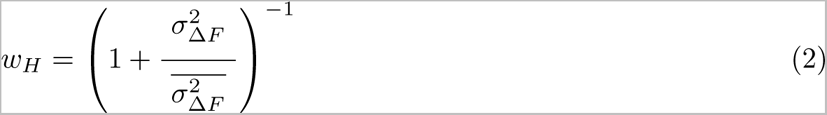

To emphasize the high-resolution features of the difference maps, we excluded low resolution reflections (*>* 5.0 Å) from the maps following Schmidt *et al.* [54]. To facilitate the reproducibility of these difference maps, we added a command-line script, *rs.diffmap*, to the *rs-booster* command-line interface of *reciprocalspaceship* [45]. The maps produced in this research used the arguments: *-a 0.0*, to achieve the weight function above, and *–dmax 5.0*, to exclude low-resolution reflections.

### Validation of Temperature-resolved Difference Maps

To rule our artifacts, we used interleaved datasets collected at 295 K to assess radiation damage and reversibility of temperature-dependent effects, and further used two crystals with reversed temperature sequences to rule out hysteresis (Fig. S1A). Indeed, the refined hinge distance was reversible and did not depend on the order of temperature changes, suggesting our temperature ramps allowed sufficient equilibration time (Fig. S1B). Isomorphous difference maps between different temperatures obtained from single crystals exhibited notably stronger difference density than maps computed between datasets collected at the same temperature (Fig. S1C and S1D), confirming that the temperature difference explains the observed effects. Equivalent temperature-resolved differences from two independent crystals were strongly correlated (Fig. S1E), demonstrating reproducibility.

## Electric-field-stimulated X-ray (EF-X) Diffraction

### Experimental Apparatus and Data Collection

We conducted the EF-X experiments at BioCARS (Advanced Photon Source, Argonne National Laboratory) using an experimental apparatus based on work by Hekstra *et al.* [23], with several important modifications that reduced sample attrition. These improvements are summarized below, and will be described in detail in an upcoming publication. The electrodes in the original experiment used wires threaded within glass capillaries, which could become retracted during sample handling, damage the crystal, and result in an osmotic mismatch with the crystal. To resolve this problem, we constructed solid state electrodes with flush surfaces for crystal contact. We produced bottom electrodes by threading tungsten wire (41 µm diameter) into glass microcapillaries (0.018 in O.D., 0.0035 in I.D., 16 mm length; Drummond) and fusing the glass around the tungsten with a Bunsen burner. We trimmed the protruding wires at the melted ends of the capillaries, and polished the electrode tips using a series of fine grit sandpapers to make a flat, flush surface with an exposed conductive patch. These bottom electrodes were placed in 3D-printed inserts compatible with reusable goniobases (Mitegen, SKU: GB-B3-R-20).

In addition, the original apparatus used a top electrode with an integrated pneumatic pump to establish liquid contact with the crystal [23]. This design required brief exposure of the crystal to the air as liquid contact was being established, risking crystal dehydration. Here, we mounted crystals on the bottom electrodes and used Sylgard 184 (Dow-Corning) to insulate their electrical contact as previously described [23]; however, we also pipetted a band of well solution in a polyester (PET) sleeve (MiTeGen) with approximately 10 µL of the crystal’s mother liquor (Fig. S2A). Prior to the experiment, we cut the sleeve above the liquid band and brought the top electrode through the mother liquor, maintaining a high humidity environment for the crystal for the duration of the experiment. Using an adjustable kapton sleeve fitted to the top electrode, we created a small droplet of mother liquor at the end of the top electrode that we used to establish liquid contact with the crystal.

Finally, we used a custom, dual-polarity pulse generator from FID GmbH (Burbach, Germany) to generate high-voltage pulses for EF-X experiments. This pulse generator is available at the BioCARS 14-ID-B beamline. For the experiment presented here, we used the data collection strategy described in Hekstra *et al.* [23] with the following modifications. At each crystal orientation, we collected an X-ray diffraction image without electric field (‘Off’), a diffraction image 200 ns after the application of a 250 ns high-voltage pulse at +3.5 kV, and a third image 200 ns after the application of a 250 ns pulse at -3.5 kV. We included a one second delay between images to permit crystal relaxation. After the three images at each crystal orientation, we rotated the crystal and repeated the collection sequence to fully sample reciprocal space (Fig. 4C). We collected the data reported here from 0*^◦^* to 180*^◦^*in 2*^◦^* steps, from 181*^◦^*to 361*^◦^* in 2*^◦^* steps, and from 361.5*^◦^* to 541.5*^◦^* in 1*^◦^* steps. This progression achieves rapid coverage of reciprocal space to ensure high completeness while evenly distributing the radiation dose during acquisition. The Laue X-ray pulses had a 100 ps duration and a spectrum from 1.02 *−* 1.18 Å (approximately 5% energy bandwidth), peaked at 1.04 Å.

### Data Reduction and Analysis of Reciprocal Space Signal

We indexed, refined the experimental geometry, and integrated the diffraction data using Precognition (Renz Research, Inc.). To scale and merge the time-resolved datasets while enforcing a common relative scale, we used `careless`, which employs approximate Bayesian inference to learn a generative model for the observed intensities and posterior estimates of the desired structure factor amplitudes [46]. We provided the image numbers, inferred wavelength of each observation, observed Miller indices, the interplanar spacing, and the observed spot centroid on the detector to `careless` as metadata. We chose a Student’s *t*-distribution with *ν* = 32 for the likelihood function based on the evaluation of values of *ν* in the merging of the ‘Off’ dataset in *P* 2_1_2_1_2_1_. For processing with `careless`, we provided the ‘Off’ data in both *P* 2_1_2_1_2_1_ and the electric-field-reduced-symmetry spacegroup, *P* 2_1_ and provided the +3.5 kV and -3.5 kV datasets in *P* 2_1_. Data collection and processing statistics for this EF-X dataset are presented in Table S10.

To evaluate the presence of electric-field-dependent structural changes in the time-resolved dataset, we took advantage of the crystallographic symmetry operations that were broken by the electric field. In particular, the two-fold screw axes along the *a*- and *c*-axes are broken, whereas the two-fold screw axis along the *b*-axis is preserved due to the alignment of the crystal relative to the applied electric field. We can compare the merged structure factor amplitudes between regions of reciprocal space that were formerly related by crystallographic symmetry in order to identify electric-field-dependent signal. In the ‘Off’ data, processed in *P* 2_1_, this symmetry should be intact, resulting in a half-dataset correlation coefficient of zero for the differences between the regions of reciprocal space. On the other hand, these differences should be measurable and reproducible for the datasets collected in the presence of an applied electric field, yielding a positive correlation coefficient. This metric, *CC_sym_*, is analogous to the half-dataset anomalous correlation coefficients (*CC_anom_*) used to evaluate anomalous signal, but measures breaking of a spacegroup symmetry operation, here 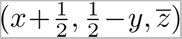, rather than Friedel’s law 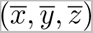. We implemented *CC_sym_* using reciprocalspaceship [45] and the result is shown in Fig. S2B.

### Extrapolated Structure Factor Refinement

To refine the excited state structure induced by the application of an electric field, we used extrapolated structure factor (ESF) refinement [23, 32]. To maximize the signal for our analysis, we refined the difference between the +3.5 kV and the -3.5 kV timepoints (‘On’ state) as follows:

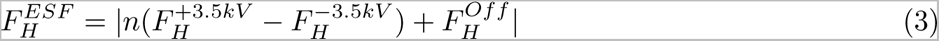

where *n* is the extrapolation factor, 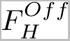 are the ‘Off’ state’s structure factor amplitudes, merged in *P* 2_1_, 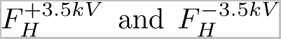 are the structure factor amplitudes for the +3.5 kV and -3.5 kV HV pulses, respectively. We scaled the 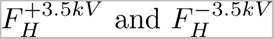 datasets relative to the 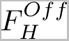 using *SCALEIT* [55], prior to computing ESFs. We computed 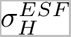 by propagating uncertainties in quadrature, and we took the absolute value of the extrapolated structure factors to avoid negative values during refinement. This assumes that the corresponding phase for the structure factor is flipped by 180*^◦^*. For refinement of the excited states, we constructed an appropriate reduced-symmetry space group by removing any crystallographic symmetry axes not collinear with the electric-field [23]. In our experiment, the crystal was mounted with the *b*-crystallographic axis offset by 24.1*±*0.5*^◦^* (mean *±* std; *N* = 1089 images) relative to the electric field vector, such that the field component along the *b* axis equals *cos*(24.1*^◦^*) *≈* 91% of the full field. In this approximation we can treat the unit cell as consisting of two copies of a redefined asymmetric unit in the *P* 1 2_1_ 1 spacegroup. To determine the extrapolation factor, we scanned values between 0 and 15 and ran automated structure refinement beginning from a model refined to the ‘Off’ data in *P* 2_1_2_1_2_1_. We found that the two copies of DHFR in the asymmetric unit refined to different hinge distances as a function of increasing *n* (Fig. S2C). The difference in hinge distance increased linearly until *n* = 8 and then plateaued at a difference of approximately 0.2 Å. As in Hekstra *et al.*, we chose the extrapolation factor to compromise between map quality (best at lower *n*) and the appearance of map features that correspond to strong peaks in the difference maps (stronger features at higher *n*) [23]. We chose an extrapolation factor of *n* = 8 for further ESF refinement because it was the lowest value (best map quality) at which the full difference in hinge distance between the two copies was realized. We used `phenix.refine` for ESF refinement [50] using isotropic B factors, occupancies, and reciprocal space-based refinement of coordinates. The refinement statistics for the ‘Off’ state from Laue diffraction and the ESF refinement of the ‘On’ state are presented in Table S11. Although ESF refinement yields higher refinement R-factors than expected for a model at 1.70 Å resolution, the magnitude of these R-factors is not a reliable measure of model quality because of the increased influence of measurement error in the extrapolated structure factors. However, since the measurement error is unchanged during refinement, relative changes in *R_work_* and *R_free_* are still useful to guide structure refinement [23]. To validate that the observed structural differences between the protein molecules of the excited-state ASU could not be explained by modeling bias, we generated simulated annealing (SA; `annealing type=cartesian`) composite omit maps using default settings in PHENIX [56, 57]. The SA composite omit maps are presented in Figure S3.

### Molecular Dynamics (MD) Simulations

To directly validate mechanistic models of the dynamics observed by X-ray diffraction, we used MD simulations of DHFR in the crystal lattice and in solvated systems. These simulations were run using *OpenMM* [58], using a custom library written to support these types of simulations (https://github.com/JBGreisman/mdtools). We ran all simulations, unless otherwise noted, in an NPT ensemble at 298 K with a 2 fs timestep, and used the Amber14SB forcefield for the protein and ions [59] and the TIP3P model for water [60]. We parameterized Folate and dihydrofolate (with and without protonation on the N5 nitrogen) using the general amber forcefield (GAFF) [61] and obtained amber-compatible NADP^+^ and NADPH parameters from the Bryce group’s database of cofactors (http://amber.manchester.ac.uk) [62, 63]. We used a native SAD structure of DHFR:NADP^+^:FOL, PDB: 7LVC, as the starting model [43], which was prepared by removing alternate conformations and protonating ionizable groups consistently with their local environments. We ran initial simulations in a 65 Å^3^ waterbox, with 200 mM NaCl. We ran 20 independent simulations that included 10 ns of equilibration followed by 500 ns production runs, outputting frames every 250 ps. We analyzed the resulting trajectories using *MDTraj* [64].

### MD Simulations of a DHFR Crystal

To simulate DHFR in its crystal context, we applied the *P* 2_1_2_1_2_1_ symmetry operations to the 7LVC starting model to build up the unit cell. We built a 3 *×* 2 *×* 1 supercell by repeating the unit cell three times along the *a* axis and twice along the *b* axis. An important consideration for such simulations is the amount of water needed to maintain the crystallographic volume. We determined this using NPT ”squeeze” runs, in which waters are added to the simulation box and strong distance restraints are slowly tapered off. More waters are then added or removed until the desired box volume is maintained within a user-determined tolerance [65]. We automated this protocol in `mdtools` and used it to generate a 3 *×* 2 *×* 1 DHFR supercell within 0.05% of the experimental volume. Additionally, we added chloride ions to the simulation box to neutralize the excess positive charge from the crystallographically observed manganese ions [43], which were included in these simulations. To equilibrate the system, we ran 50 ns of MD in an NPT ensemble. We then initialized production simulations in an NVT ensemble from the last frame of equilibration. We ran three independent production simulations for 500 ns, outputting frames every 100 ps.

### Classification of Met20 loop substates in simulation

We quantified the population of the two Met20 loop substates using the Trp22-*ϕ* dihedral as a reporter. Since this dihedral exhibited two distinct states, we fit the data to a two-state Gaussian mixture model using all frames from each trajectory. We used the Gaussian mixture model implemented in *scikit-learn* for this analysis [66]. To estimate the uncertainty in this classification, we classified the frames of each trajectory independently using the fit model and reported the mean and standard error across the trajectories. This analysis was repeated for the simulations of the solvated and lattice systems. For the solvated system, we used twenty independent trajectories to quantify the population of each substate. For the lattice system, we treated each protein molecules in the simulation independently, yielding 72 independent trajectories (24 protein molecules *×* 3 simulations).

### Biased MD Simulations in Bulk Solvent

To validate that the results observed from X-ray diffraction experiments are recapitulated outside of the crystal context we ran MD simulations of the model of the DHFR Michaelis complex, using the same solvated simulation system as our unbiased trajectories. In order to bias the sampling of the MD simulations based on the hinge distance, we added a custom distance restraint between the *C_α_* atoms of Asn23 and Pro53 using the following functional form:

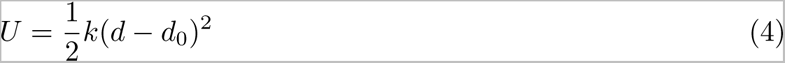

where *k* was chosen to be 50.0 kcal/mol/Å^2^, *d* is the distance between the *C_α_* atoms of Asn23 and Pro53 under the minimum periodic image convention, and *d*_0_ is the desired equilibrium distance for the active site cleft. We ran MD simulations with *d*_0_ values of 18.8, 19.2, 19.6, 20.0, and 20.4 Å in order to bias the sampling across the range of crystallographically observed values. 100 independent simulations were equilibrated for 10 ns and then simulated for 100 ns for each value of *d*_0_.

### MD Simulations of the Reactive Ternary Complex in Bulk Solvent

Using the 7LVC starting model, we modeled NADPH and dihydrofolate (protonated and deprotonated) to represent the reactive ternary complex of DHFR. We prepared the simulation system in a 65 Å^3^ waterbox with 200 mM of NaCl, and we ran 50 independent simulations with 10 ns of equilibration and then 100 ns production simulations.

## Supplementary Figures and Tables

**Figure S1:**
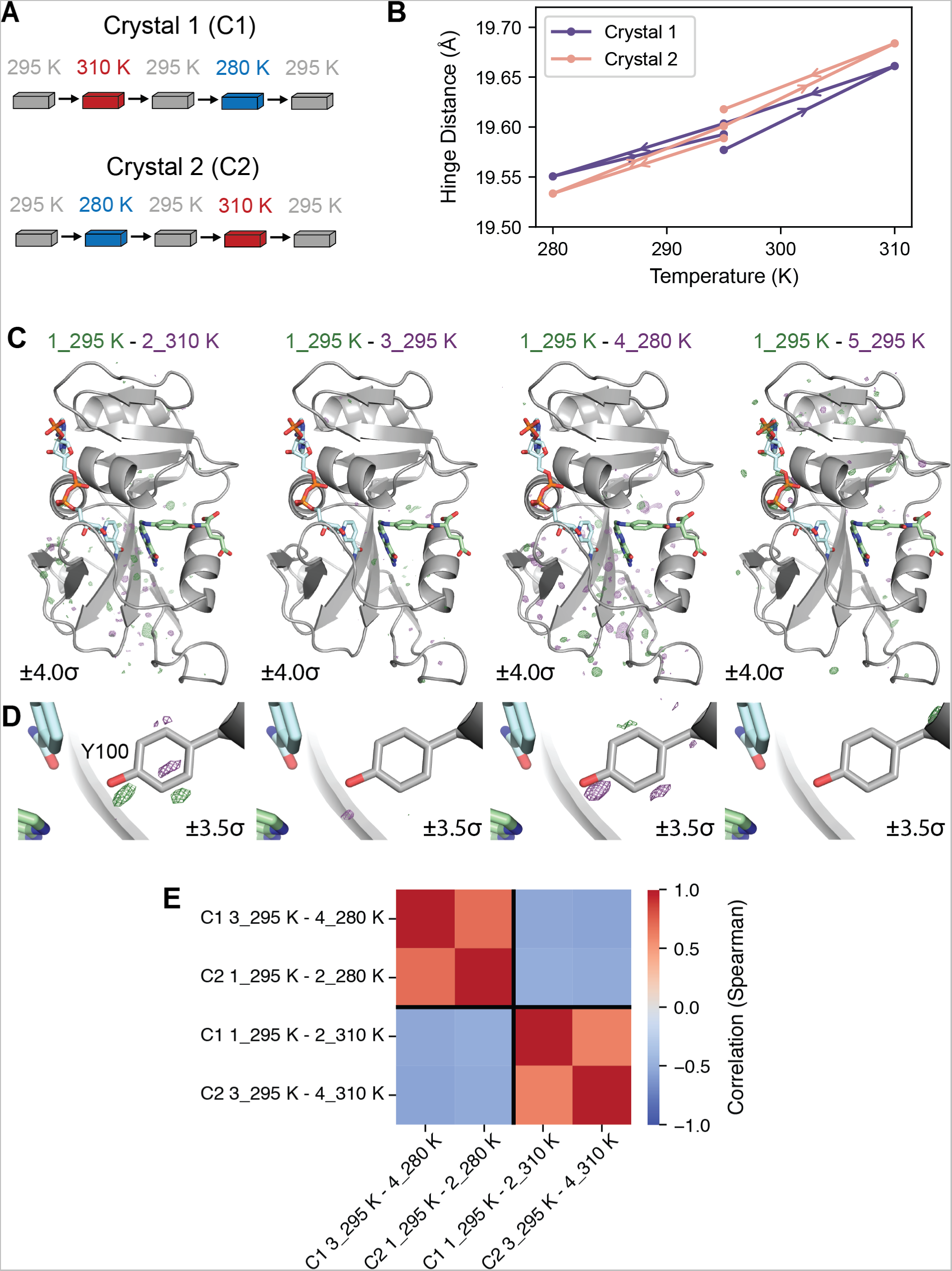
Reversibility and reproducibility of multi-temperature diffraction experiments. (A) Schematic of single-crystal, multi-temperature diffraction experiments. (B) Plots of the refined hinge distance versus temperature for both single-crystal experiments demonstrate that the experiment is reversible. (C) Temperature-resolved difference maps between the first dataset from crystal 1 and the subsequent four datasets. More significant density peaks are observed for maps generated from datasets collected at different temperatures. (D) Zoom-in on Tyr100 in the difference maps emphasizes that observed features are temperature-dependent (carved within 2 Å of Tyr100). (E) Heatmap of the Spearman correlation coefficients between difference structure factor amplitudes computed from independent single-crystal experiments. Equivalent temperature changes yield strongly correlated difference amplitudes, while the opposite temperature changes produce strongly anti-correlated results. This demonstrates that the observed structural changes in the single-crystal, multi-temperature experiments are reproducible between independent experiments.

**Fig. S2.**
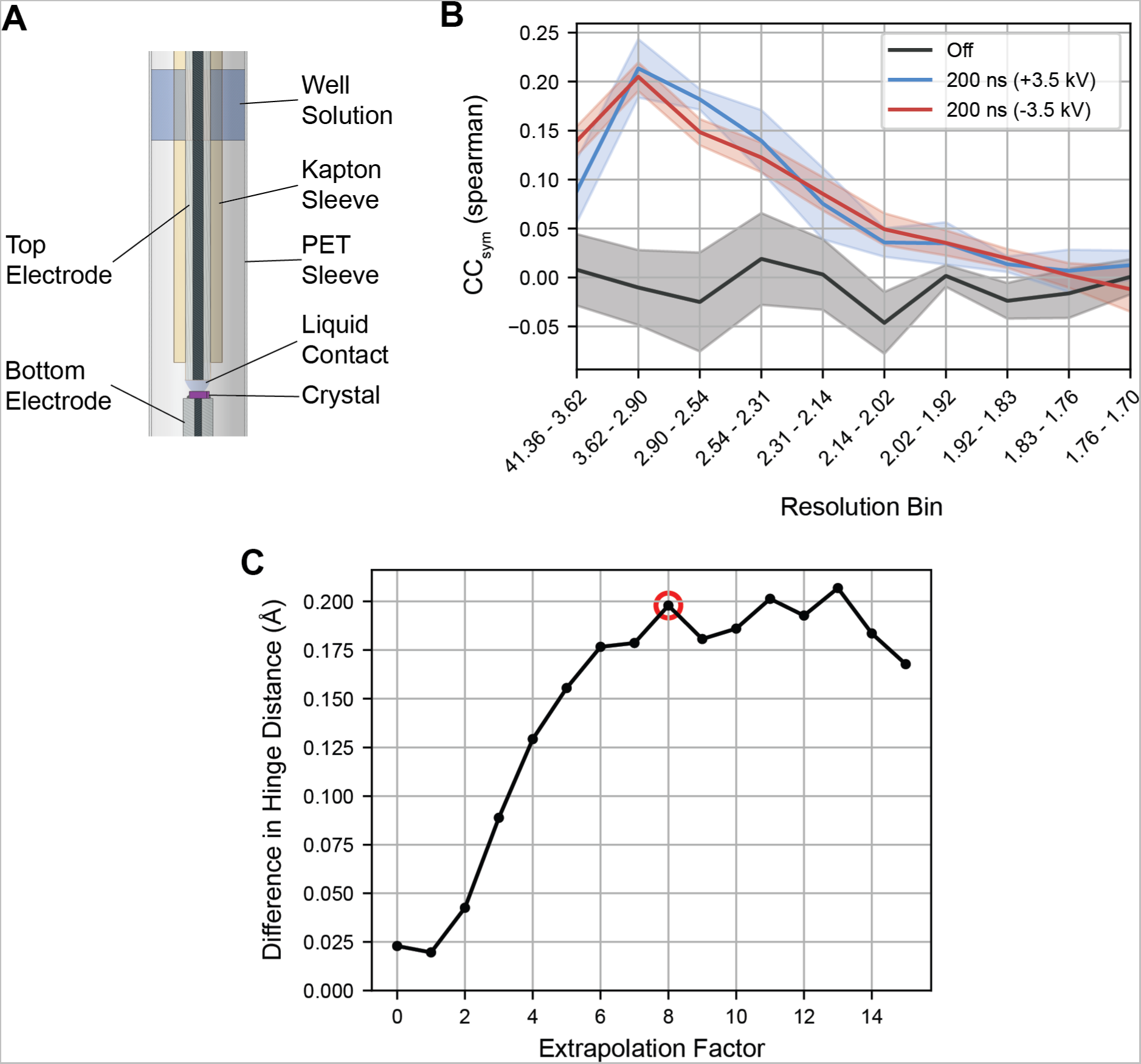
Experimental apparatus and analysis for electric-field-stimulated X-ray diffraction of *ec*DHFR. (A) Diagram of the revised experimental apparatus for EF-X. Liquid contact is made within a band of well solution that is osmotically matched to the crystal, ensuring a high humidity environment for the duration of the experiment. (B) Plot of *CCsym* versus resolution bin. *CCsym* is an indicator of the reproducibility of observed symmetry breaking during an EF-X experiment. The 95% confidence interval from 5 random partitions of the diffraction images is shown. For the ‘Off’ dataset in which the symmetry operation is preserved, no significant correlation between half-datasets is expected because differences for symmetry-related observations should only reflect experimental error. The positive correlations for differences measured during the high-voltage pulses indicates significant electric-field-dependent symmetry breaking.(C) Plot of the refined difference in hinge distance between the two copies of DHFR in the *P* 2_1_ ASU as a function of extrapolation factor. With an extrapolation factor of zero, the data is equivalent to ‘Off’ structure factor amplitudes processed in the reduced-symmetry spacegroup. The difference in hinge distance increases linearly with extrapolation factor until a value of 8 and plateaus at a difference of approximately 0.2 Å. The extrapolation factor chosen for ESF refinement of the excited state is indicated with a red circle.

**Fig. S3.**
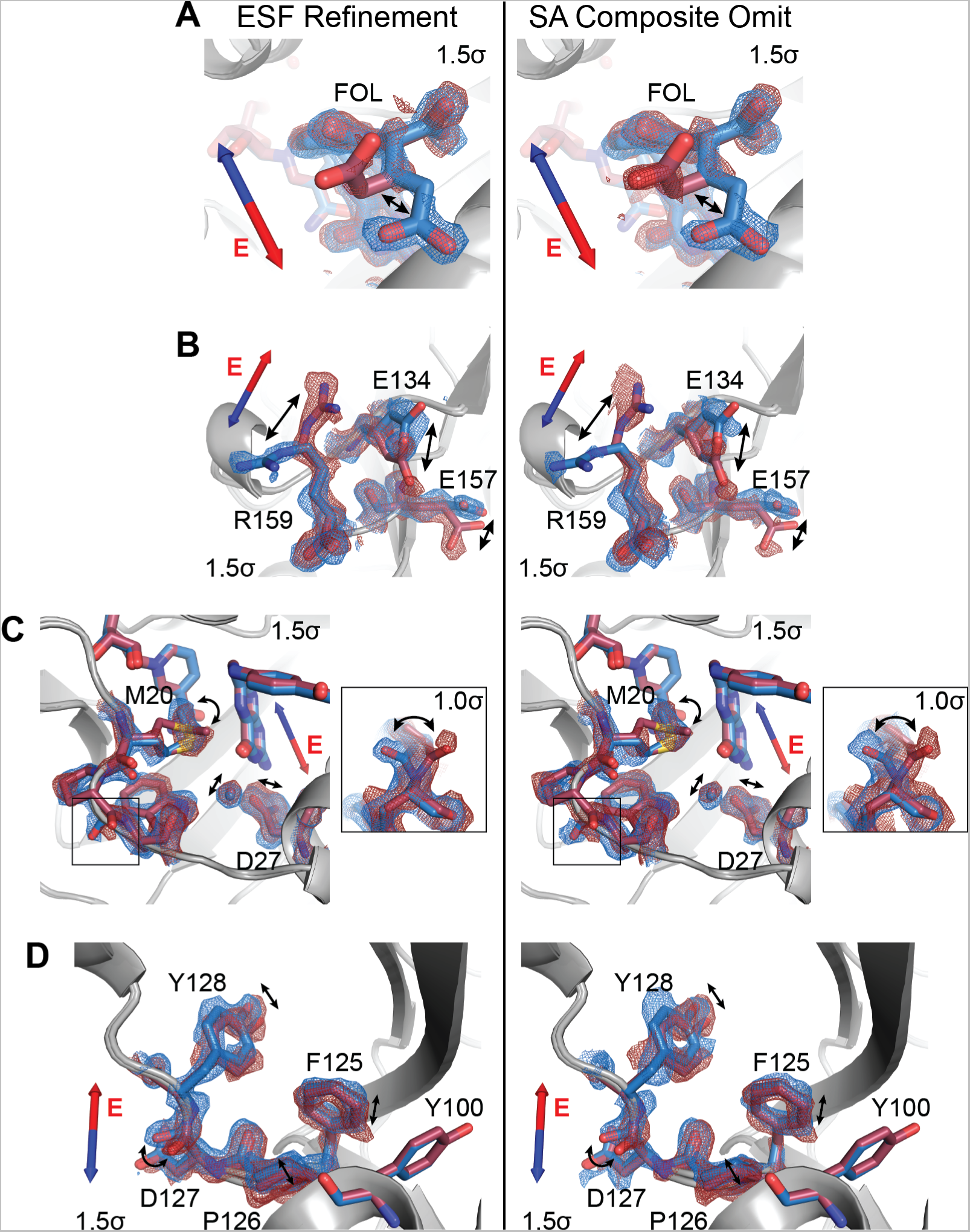
Composite omit maps validate modeling of EF-X excited state. (A) to (D) Comparison of 2*mFo − DFc* maps from ESF refinement (left column) and corresponding simulated annealing (SA) composite omit maps (right column). Superposed models and maps from both protein molecules of the excited-state ASU highlight electric-field induced structural changes. Blue and red arrows depict electric field vector for the blue and red models, respectively, and maps are contoured at 1.5*σ* and carved within 1.5 Å of shown atoms. (A) Carboxylate sidechain of folate and (B) charged sidechains near the C-terminus demonstrate electric-field-dependent structural changes consistent with the formal charges of the residues. (C) Active site residues and Pro21 backbone carbonyl (inset; contoured at 1.0*σ*) differs between protein molecules. (D) Conformational changes among residues 125 to 128. The similarity between the electron density maps from ESF refinement and the SA composite omit maps indicates that the observed structural differences between the molecules of the excited-state ASU are not the result of modeling bias.

**Fig. S4.**
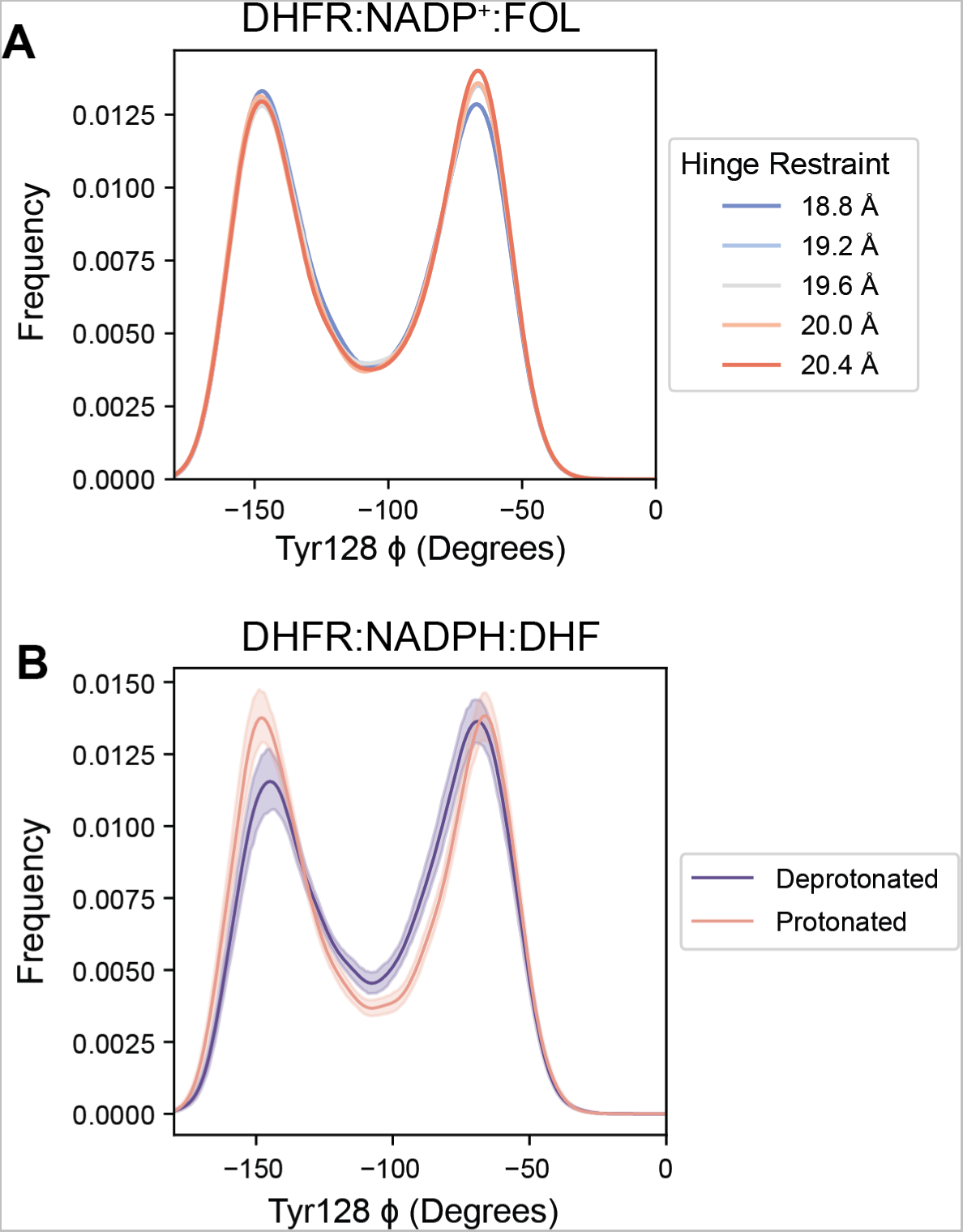
Tyr128 backbone conformations in MD simulations. (A) Kernel density estimates of the Tyr128-*ϕ* dihedral from MD simulations at each imposed hinge distance restraint. The Tyr128-*ϕ* dihedral does not exhibit a monotonic relationship as a function of hinge distance. (B) Kernel density estimates of the Tyr128-*ϕ* dihedral from MD simulations of the reactive ternary complex (95% confidence interval is shown). The Tyr128-*ϕ* dihedral distribution is altered by substrate protonation.

**Table S1.** Summary statistics for DHFR:NADP^+^:MFOL complex

|  |  |
| --- | --- |
| PDB ID | 8DAI |
| Temperature | 285 K |
| Data Collection<sup>1</sup> |  |
| Wavelength (Å) | 0.9537 |
| Spacegroup | *P*2<sub>1</sub>2<sub>1</sub>2<sub>1</sub> |
| Cell dimensions (Å) |  |
| a, b, c | 34.25, 45.36, 98.85 |
| Total observations | 2,736,784 |
| Unique observations | 105,471 |
| Resolution (Å) | 49.42 - 1.14<br>(1.16 - 1.14) |
| Multiplicity | 25.9 (14.4) |
| Completeness (%) | 97.2 (73.0) |
| Mean *I*/σ<sub>*I*</sub> | 11.9 (0.4) |
| R<sub>pim</sub> | 0.028 (0.980) |
| CC<sub>1/2</sub> | 0.999 (0.326) |
| Refinement<sup>2</sup> |  |
| R<sub>work</sub> (%) | 12.68 |
| R<sub>free</sub> (%) | 16.00 |
| R.M.S. Deviations |  |
| Bonds (Å) | 0.013 |
| Angles (°) | 1.357 |
| Wilson B (Å<sup>2</sup>) | 15.57 |
| Mean B factor (Å<sup>2</sup>) |  |
| Total | 22.97 |
| Macromolecules | 21.07 |
| Ligands | 21.71 |
| Water | 39.71 |
| Clashscore | 2.23 |
| Ramachandran |  |
| Favored (%) | 98.70 |
| Allowed (%) | 1.30 |
| Outliers (%) | 0.00 |

<sup>1</sup> Reported by *dials.scale* in *DIALS* [44]
<sup>2</sup> Reported by *PHENIX* [50]

**Table S2.**
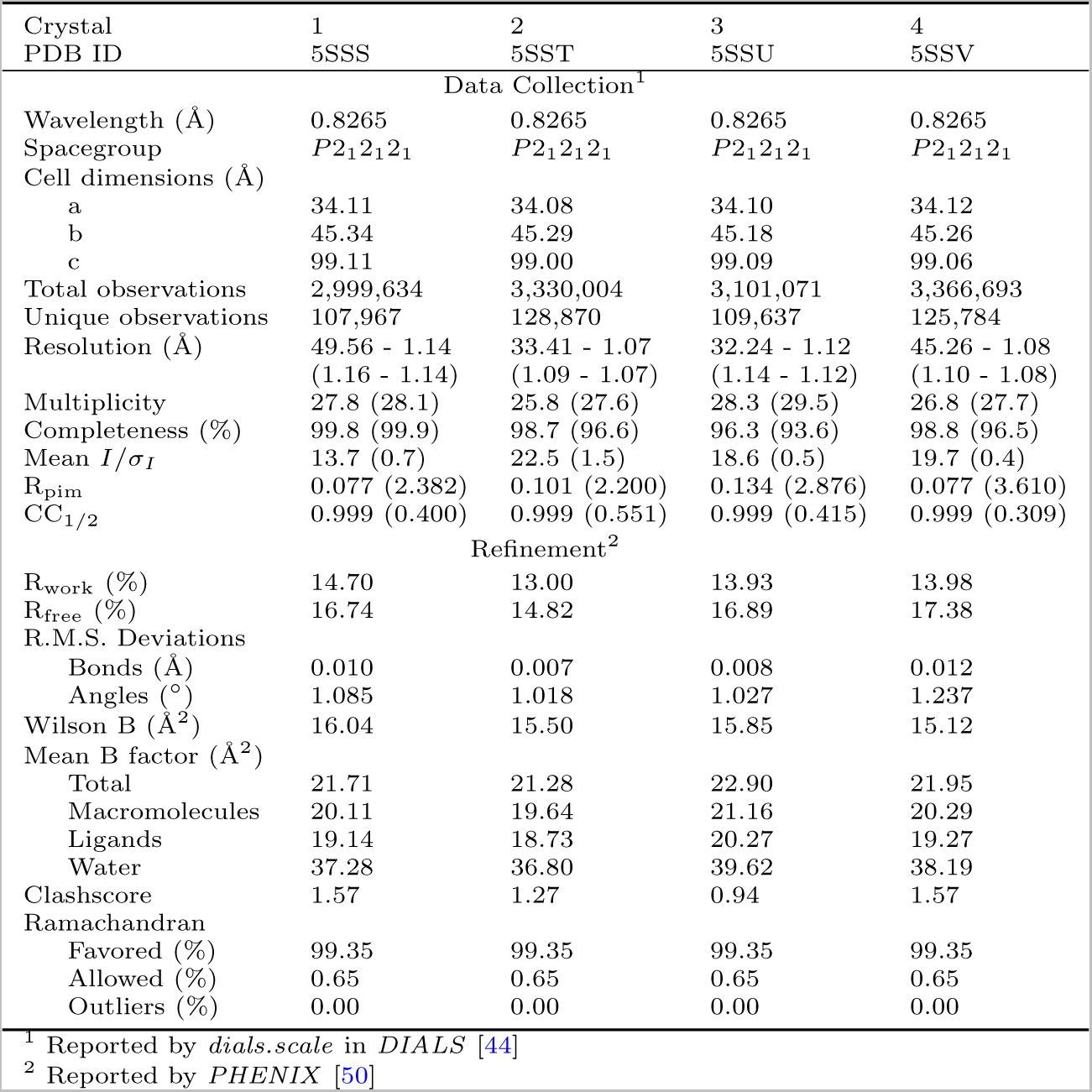
Summary statistics for datasets at 270 K

**Table S3.**
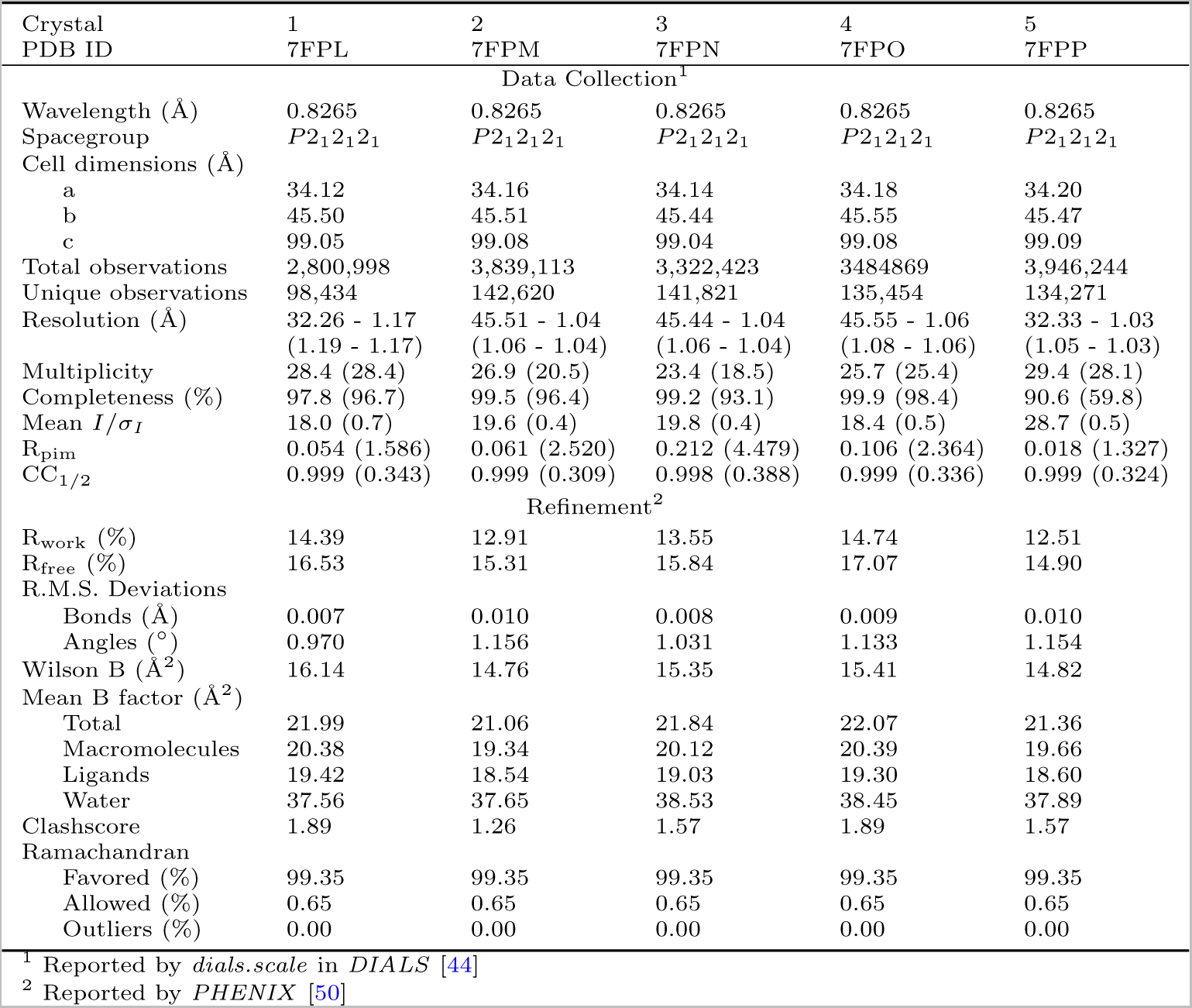
Summary statistics for datasets at 280 K

**Table S4.** Summary statistics for datasets at 290 K

| Crystal<br>PDB ID | 1<br>7FPR | 2<br>7FPS | 3<br>7FPT | 4<br>7FPU | 5<br>7FPV |
| --- | --- | --- | --- | --- | --- |
|  | Data Collection<sup>1</sup> |  |  |  |  |
| Wavelength (Å) | 0.8265 | 0.8265 | 0.8265 | 0.8265 | 0.8265 |
| Spacegroup | *P*2<sub>1</sub>2<sub>1</sub>2<sub>1</sub> | *P*2<sub>1</sub>2<sub>1</sub>2<sub>1</sub> | *P*2<sub>1</sub>2<sub>1</sub>2<sub>1</sub> | *P*2<sub>1</sub>2<sub>1</sub>2<sub>1</sub> | *P*2<sub>1</sub>2<sub>1</sub>2<sub>1</sub> |
| Cell dimensions (Å) |  |  |  |  |  |
| a | 34.18 | 34.19 | 34.19 | 34.18 | 34.20 |
| b | 45.49 | 45.56 | 45.59 | 45.60 | 45.56 |
| c | 99.10 | 99.05 | 99.05 | 99.07 | 99.04 |
| Total observations | 3,627,079 | 2,149,046 | 3,640,008 | 3,834,528 | 3,765,323 |
| Unique observations | 125,984 | 80,476 | 123,429 | 132,517 | 142,372 |
| Resolution (Å) | 32.31 - 1.07<br>(1.09 - 1.07) | 49.53 - 1.26<br>(1.28 - 1.26) | 32.32 - 1.07<br>(1.09 - 1.07) | 32.31 - 1.05<br>(1.07 - 1.05) | 41.39 - 1.04<br>(1.06 - 1.04) |
| Multiplicity | 28.8 (29.4) | 26.7 (23.3) | 29.5 (30.2) | 28.9 (27.8) | 26.4 (20.7) |
| Completeness (%) | 95.6 (92.8) | 99.8 (98.4) | 93.5 (90.4) | 94.8 (80.5) | 99.2 (95.4) |
| Mean *I*/$\sigma_I$ | 23.7 (0.6) | 18.6 (1.1) | 25.8 (0.8) | 24.5 (0.6) | 27.8 (0.5) |
| R<sub>pim</sub> | 0.111 (2.284) | 0.105 (1.704) | 0.028 (1.401) | 0.027 (1.713) | 0.031 (1.101) |
| CC<sub>1/2</sub> | 0.999 (0.312) | 0.999 (0.357) | 0.999 (0.555) | 0.999 (0.349) | 0.999 (0.364) |
|  | Refinement<sup>2</sup> |  |  |  |  |
| R<sub>work</sub> (%) | 12.37 | 13.17 | 11.84 | 12.25 | 12.60 |
| R<sub>free</sub> (%) | 14.62 | 16.65 | 14.22 | 14.72 | 14.50 |
| R.M.S. Deviations |  |  |  |  |  |
| Bonds (Å) | 0.008 | 0.009 | 0.009 | 0.009 | 0.007 |
| Angles (°) | 1.035 | 1.083 | 1.144 | 1.124 | 1.025 |
| Wilson B (Å<sup>2</sup>) | 14.98 | 15.94 | 15.00 | 15.29 | 14.11 |
| Mean B factor (Å<sup>2</sup>) |  |  |  |  |  |
| Total | 23.05 | 22.82 | 21.68 | 21.81 | 21.80 |
| Macromolecules | 21.04 | 20.95 | 19.93 | 20.10 | 19.84 |
| Ligands | 19.85 | 20.08 | 18.66 | 18.81 | 18.35 |
| Water | 42.52 | 40.85 | 38.81 | 38.55 | 41.08 |
| Clashscore | 1.26 | 2.20 | 1.57 | 1.57 | 1.89 |
| Ramachandran |  |  |  |  |  |
| Favored (%) | 99.35 | 99.35 | 99.35 | 99.35 | 99.35 |
| Allowed (%) | 0.65 | 0.65 | 0.65 | 0.65 | 0.65 |
| Outliers (%) | 0.00 | 0.00 | 0.00 | 0.00 | 0.00 |

<sup>1</sup> Reported by *dials.scale* in *DIALS* [44]<sup>2</sup> Reported by *PHENIX* [50]

**Table S5.**
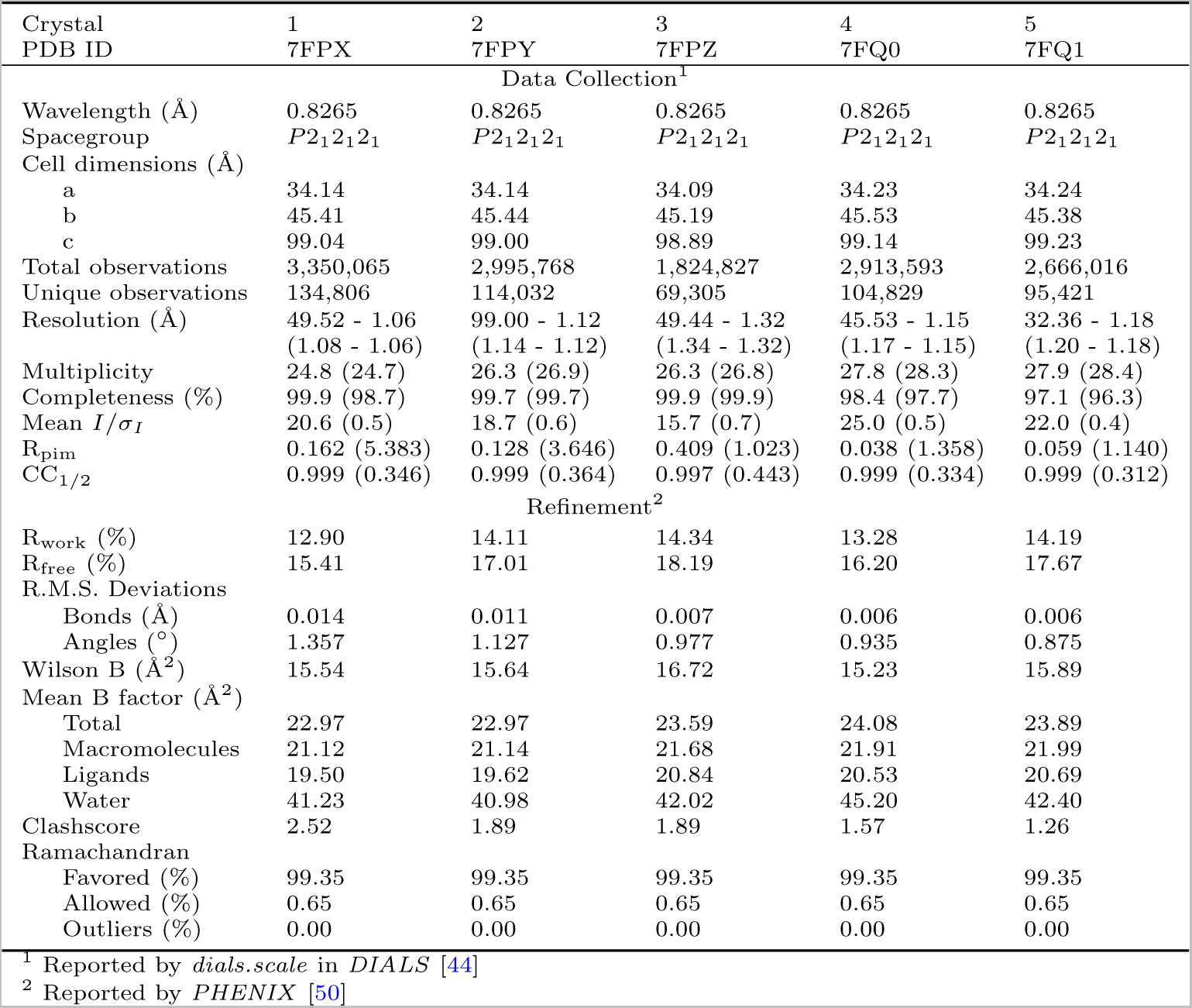
Summary statistics for datasets at 300 K

**Table S6.** Summary statistics for datasets at 310 K

| Crystal<br>PDB ID | 1<br>7FQ3 | 2<br>7FQ4 | 3<br>7FQ5 |
| --- | --- | --- | --- |
|  | Data Collection<sup>1</sup> |  |  |
| Wavelength (Å) | 0.8265 | 0.8265 | 0.8265 |
| Spacegroup | *P*2<sub>1</sub>2<sub>1</sub>2<sub>1</sub> | *P*2<sub>1</sub>2<sub>1</sub>2<sub>1</sub> | *P*2<sub>1</sub>2<sub>1</sub>2<sub>1</sub> |
| Cell dimensions (Å) |  |  |  |
| a | 34.18 | 34.15 | 34.19 |
| b | 45.49 | 45.23 | 45.30 |
| c | 99.33 | 99.22 | 99.25 |
| Total observations | 1,969,232 | 1,829,107 | 1,788,359 |
| Unique observations | 73,267 | 67,810 | 65,420 |
| Resolution (Å) | 41.36 - 1.30<br>(1.32 - 1.30) | 33.42 - 1.33<br>(1.36 - 1.33) | 99.25 - 1.35<br>(1.37 - 1.35) |
| Multiplicity | 26.9 (27.1) | 27.3 (28.6) | 27.3 (28.9) |
| Completeness (%) | 99.4 (92.4) | 99.1 (97.9) | 99.5 (89.5) |
| Mean *I*/$\sigma_I$ | 15.4 (0.4) | 24.5 (0.6) | 21.3 (0.5) |
| R<sub>pim</sub> | 0.310 (1.596) | 0.080 (0.848) | 0.108 (1.048) |
| CC<sub>1/2</sub> | 0.997 (0.360) | 0.999 (0.301) | 0.999 (0.328) |
|  | Refinement<sup>2</sup> |  |  |
| R<sub>work</sub> (%) | 15.24 | 14.61 | 14.96 |
| R<sub>free</sub> (%) | 18.66 | 18.57 | 18.96 |
| R.M.S. Deviations |  |  |  |
| Bonds (Å) | 0.006 | 0.008 | 0.009 |
| Angles (°) | 1.001 | 1.037 | 1.007 |
| Wilson B (Å<sup>2</sup>) | 17.48 | 15.77 | 17.55 |
| Mean B factor (Å<sup>2</sup>) |  |  |  |
| Total | 23.71 | 24.37 | 25.03 |
| Macromolecules | 21.58 | 22.22 | 22.77 |
| Ligands | 20.69 | 21.76 | 21.86 |
| Water | 44.24 | 44.80 | 46.82 |
| Clashscore | 1.89 | 2.20 | 3.15 |
| Ramachandran |  |  |  |
| Favored (%) | 99.35 | 99.35 | 99.35 |
| Allowed (%) | 0.65 | 0.65 | 0.65 |
| Outliers (%) | 0.00 | 0.00 | 0.00 |

<sup>1</sup> Reported by *dials.scale* in *DIALS* [44]<sup>2</sup> Reported by *PHENIX* [50]

**Table S7.**
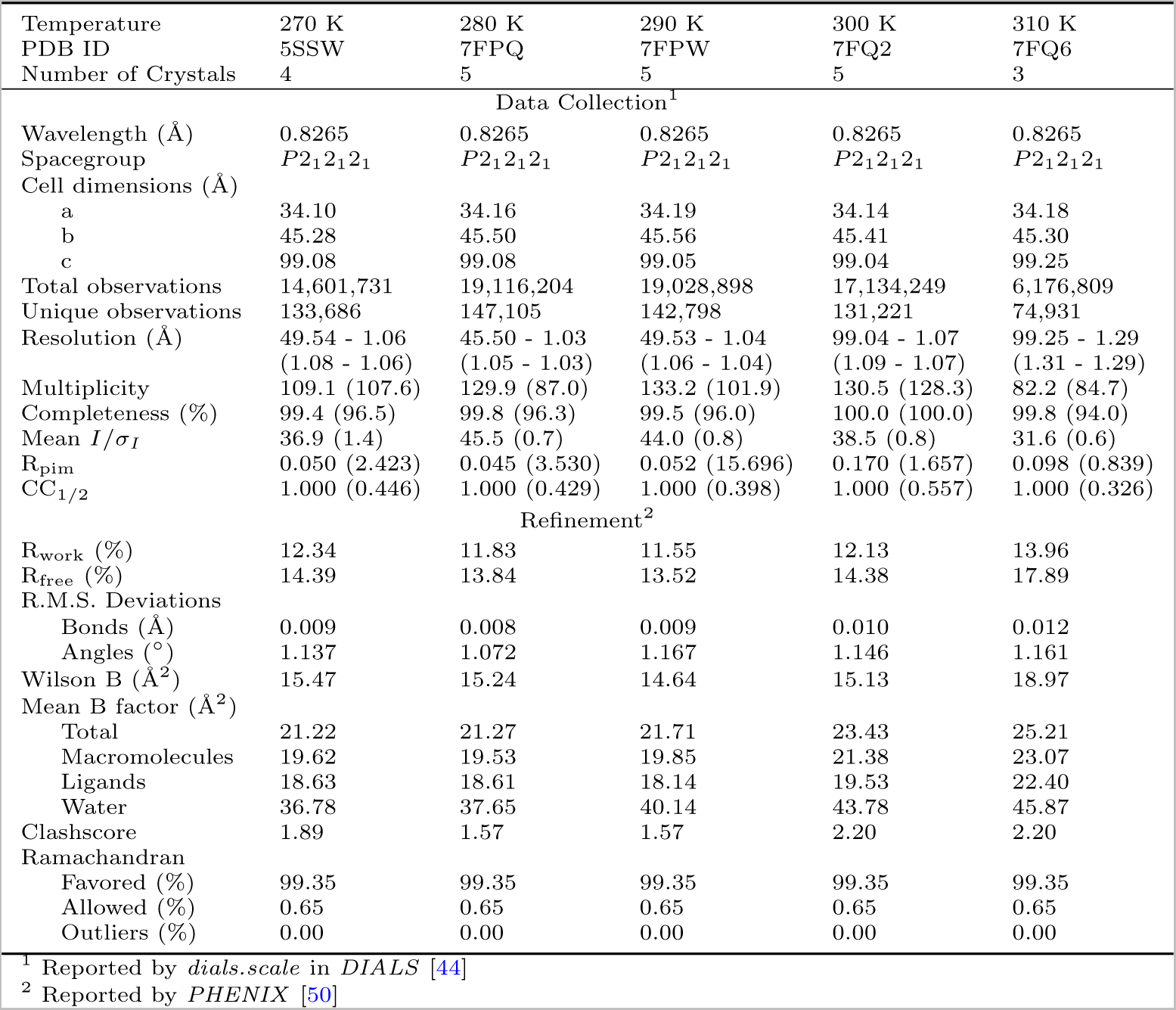
Summary statistics for multi-crystal, multi-temperature datasets

| Temperature | 270 K | 280 K | 290 K | 300 K | 310 K |
| --- | --- | --- | --- | --- | --- |
| PDB ID | 5SSW | 7FPQ | 7FPW | 7FQ2 | 7FQ6 |
| Number of Crystals | 4 | 5 | 5 | 5 | 3 |
| Data Collection<sup>1</sup> |  |  |  |  |  |
| Wavelength (Å) | 0.8265 | 0.8265 | 0.8265 | 0.8265 | 0.8265 |
| Spacegroup | $P2_12_12_1$ | $P2_12_12_1$ | $P2_12_12_1$ | $P2_12_12_1$ | $P2_12_12_1$ |
| Cell dimensions (Å) |  |  |  |  |  |
| a | 34.10 | 34.16 | 34.19 | 34.14 | 34.18 |
| b | 45.28 | 45.50 | 45.56 | 45.41 | 45.30 |
| c | 99.08 | 99.08 | 99.05 | 99.04 | 99.25 |
| Total observations | 14,601,731 | 19,116,204 | 19,028,898 | 17,134,249 | 6,176,809 |
| Unique observations | 133,686 | 147,105 | 142,798 | 131,221 | 74,931 |
| Resolution (Å) | 49.54 - 1.06 | 45.50 - 1.03 | 49.53 - 1.04 | 99.04 - 1.07 | 99.25 - 1.29 |
|  | (1.08 - 1.06) | (1.05 - 1.03) | (1.06 - 1.04) | (1.09 - 1.07) | (1.31 - 1.29) |
| Multiplicity | 109.1 (107.6) | 129.9 (87.0) | 133.2 (101.9) | 130.5 (128.3) | 82.2 (84.7) |
| Completeness (%) | 99.4 (96.5) | 99.8 (96.3) | 99.5 (96.0) | 100.0 (100.0) | 99.8 (94.0) |
| Mean $I/\sigma_I$ | 36.9 (1.4) | 45.5 (0.7) | 44.0 (0.8) | 38.5 (0.8) | 31.6 (0.6) |
| R<sub>pim</sub> | 0.050 (2.423) | 0.045 (3.530) | 0.052 (15.696) | 0.170 (1.657) | 0.098 (0.839) |
| CC<sub>1/2</sub> | 1.000 (0.446) | 1.000 (0.429) | 1.000 (0.398) | 1.000 (0.557) | 1.000 (0.326) |
| Refinement<sup>2</sup> |  |  |  |  |  |
| R<sub>work</sub> (%) | 12.34 | 11.83 | 11.55 | 12.13 | 13.96 |
| R<sub>free</sub> (%) | 14.39 | 13.84 | 13.52 | 14.38 | 17.89 |
| R.M.S. Deviations |  |  |  |  |  |
| Bonds (Å) | 0.009 | 0.008 | 0.009 | 0.010 | 0.012 |
| Angles (°) | 1.137 | 1.072 | 1.167 | 1.146 | 1.161 |
| Wilson B (Å<sup>2</sup>) | 15.47 | 15.24 | 14.64 | 15.13 | 18.97 |
| Mean B factor (Å<sup>2</sup>) |  |  |  |  |  |
| Total | 21.22 | 21.27 | 21.71 | 23.43 | 25.21 |
| Macromolecules | 19.62 | 19.53 | 19.85 | 21.38 | 23.07 |
| Ligands | 18.63 | 18.61 | 18.14 | 19.53 | 22.40 |
| Water | 36.78 | 37.65 | 40.14 | 43.78 | 45.87 |
| Clashscore | 1.89 | 1.57 | 1.57 | 2.20 | 2.20 |
| Ramachandran |  |  |  |  |  |
| Favored (%) | 99.35 | 99.35 | 99.35 | 99.35 | 99.35 |
| Allowed (%) | 0.65 | 0.65 | 0.65 | 0.65 | 0.65 |
| Outliers (%) | 0.00 | 0.00 | 0.00 | 0.00 | 0.00 |

<sup>1</sup> Reported by *dials.scale* in *DIALS* [44]<sup>2</sup> Reported by *PHENIX* [50]

**Table S8.** Summary statistics for single-crystal, multi-temperature datasets (crystal 1)

| Temperature | 295 K | 310 K | 295 K | 280 K | 295 K |
| --- | --- | --- | --- | --- | --- |
| Pass on Crystal | 1 | 2 | 3 | 4 | 5 |
| PDB ID | 7FQ7 | 7FQ8 | 7FQ9 | 7FQA | 7FQB |
|  | Data Collection<sup>1</sup> |  |  |  |  |
| Wavelength (Å) | 0.8265 | 0.8265 | 0.8265 | 0.8265 | 0.8265 |
| Spacegroup | $P2_12_12_1$ | $P2_12_12_1$ | $P2_12_12_1$ | $P2_12_12_1$ | $P2_12_12_1$ |
| Cell dimensions (Å) |  |  |  |  |  |
| a | 34.26 | 34.29 | 34.27 | 34.20 | 34.22 |
| b | 45.59 | 45.65 | 45.63 | 45.46 | 45.47 |
| c | 98.96 | 99.03 | 98.97 | 98.99 | 99.02 |
| Total observations | 3,400,772 | 2,601,689 | 3,315,426 | 3,572,742 | 3,053,872 |
| Unique observations | 117,634 | 89,462 | 115,024 | 123,443 | 105,250 |
| Resolution (Å) | 32.37 - 1.10 | 41.46 - 1.21 | 41.44 - 1.11 | 32.32 - 1.08 | 32.34 - 1.14 |
|  | (1.12 - 1.10) | (1.23 - 1.21) | (1.13 - 1.11) | (1.10 - 1.08) | (1.16 - 1.14) |
| Multiplicity | 28.9 (28.8) | 29.1 (29.5) | 28.8 (28.6) | 28.9 (29.0) | 29.0 (28.0) |
| Completeness (%) | 96.7 (95.6) | 97.6 (96.6) | 97.1 (96.4) | 96.5 (94.5) | 96.6 (96.2) |
| Mean $I/\sigma_I$ | 12.8 (0.3) | 14.1 (0.4) | 12.6 (0.3) | 12.7 (0.3) | 13.5 (0.4) |
| R<sub>pim</sub> | 0.027 (1.213) | 0.037 (1.105) | 0.029 (1.257) | 0.025 (1.426) | 0.028 (1.245) |
| CC<sub>1/2</sub> | 0.999 (0.324) | 0.999 (0.380) | 0.999 (0.354) | 0.999 (0.311) | 0.999 (0.332) |
|  | Refinement<sup>2</sup> |  |  |  |  |
| R<sub>work</sub> (%) | 12.80 | 12.64 | 13.13 | 13.11 | 13.36 |
| R<sub>free</sub> (%) | 15.89 | 16.40 | 15.94 | 15.89 | 16.48 |
| R.M.S. Deviations |  |  |  |  |  |
| Bonds (Å) | 0.008 | 0.011 | 0.007 | 0.005 | 0.006 |
| Angles (°) | 1.054 | 1.192 | 1.008 | 0.884 | 0.953 |
| Wilson B (Å<sup>2</sup>) | 17.06 | 16.74 | 17.13 | 17.26 | 16.81 |
| Mean B factor (Å<sup>2</sup>) |  |  |  |  |  |
| Total | 22.18 | 23.86 | 23.34 | 21.75 | 22.30 |
| Macromolecules | 20.38 | 21.68 | 21.54 | 20.09 | 20.46 |
| Ligands | 18.80 | 19.14 | 19.88 | 18.77 | 18.97 |
| Water | 39.94 | 45.87 | 41.17 | 38.11 | 40.38 |
| Clashscore | 1.26 | 2.20 | 1.26 | 1.26 | 1.89 |
| Ramachandran |  |  |  |  |  |
| Favored (%) | 99.35 | 99.35 | 99.35 | 99.35 | 99.35 |
| Allowed (%) | 0.65 | 0.65 | 0.65 | 0.65 | 0.65 |
| Outliers (%) | 0.00 | 0.00 | 0.00 | 0.00 | 0.00 |

<sup>1</sup> Reported by *dials.scale* in *DIALS* [44]<sup>2</sup> Reported by *PHENIX* [50]

**Table S9.** Summary statistics for single-crystal, multi-temperature datasets (crystal 2)

| Temperature | 295 K | 280 K | 295 K | 310 K | 295 K |
| --- | --- | --- | --- | --- | --- |
| Pass on Crystal | 1 | 2 | 3 | 4 | 5 |
| PDB ID | 7FQC | 7FQD | 7FQE | 7FQF | 7FQG |
|  | Data Collection<sup>1</sup> |  |  |  |  |
| Wavelength (Å) | 0.9795 | 0.9795 | 0.9795 | 0.9795 | 0.9795 |
| Spacegroup | *P*2<sub>1</sub>2<sub>1</sub>2<sub>1</sub> | *P*2<sub>1</sub>2<sub>1</sub>2<sub>1</sub> | *P*2<sub>1</sub>2<sub>1</sub>2<sub>1</sub> | *P*2<sub>1</sub>2<sub>1</sub>2<sub>1</sub> | *P*2<sub>1</sub>2<sub>1</sub>2<sub>1</sub> |
| Cell dimensions (Å) |  |  |  |  |  |
| a | 34.26 | 34.20 | 34.25 | 34.30 | 34.28 |
| b | 45.63 | 45.52 | 45.60 | 45.71 | 45.68 |
| c | 99.03 | 99.06 | 99.09 | 99.12 | 99.04 |
| Total observations | 2,722,807 | 2,756,746 | 2,721,740 | 2,438,928 | 2,674,944 |
| Unique observations | 97,218 | 99,044 | 97,041 | 86,426 | 95,020 |
| Resolution (Å) | 49.52 - 1.18 | 49.53 - 1.17 | 49.55 - 1.18 | 49.56 - 1.23 | 49.52 - 1.19 |
|  | (1.20 - 1.18) | (1.19 - 1.17) | (1.20 - 1.18) | (1.25 - 1.23) | (1.21 - 1.19) |
| Multiplicity | 28.0 (23.7) | 27.8 (21.2) | 28.0 (23.8) | 28.2 (28.4) | 28.1 (26.5) |
| Completeness (%) | 98.5 (94.9) | 98.2 (91.2) | 98.4 (94.4) | 98.9 (98.2) | 98.6 (95.5) |
| Mean $I/\sigma_I$ | 13.0 (0.4) | 12.8 (0.4) | 12.6 (0.4) | 12.3 (0.3) | 12.1 (0.4) |
| R<sub>pim</sub> | 0.033 (1.029) | 0.031 (1.107) | 0.033 (1.127) | 0.044 (1.404) | 0.038 (1.220) |
| CC<sub>1/2</sub> | 0.999 (0.359) | 0.999 (0.308) | 0.999 (0.308) | 0.999 (0.321) | 0.999 (0.333) |
|  | Refinement<sup>2</sup> |  |  |  |  |
| R<sub>work</sub> (%) | 12.74 | 14.09 | 12.88 | 13.06 | 13.13 |
| R<sub>free</sub> (%) | 16.00 | 16.17 | 16.38 | 16.91 | 16.41 |
| R.M.S. Deviations |  |  |  |  |  |
| Bonds (Å) | 0.008 | 0.008 | 0.009 | 0.013 | 0.005 |
| Angles (°) | 1.028 | 1.030 | 1.114 | 1.238 | 0.905 |
| Wilson B (Å<sup>2</sup>) | 17.21 | 17.63 | 17.19 | 17.08 | 16.96 |
| Mean B factor (Å<sup>2</sup>) |  |  |  |  |  |
| Total | 22.47 | 21.53 | 22.56 | 25.06 | 22.56 |
| Macromolecules | 20.65 | 19.92 | 20.71 | 22.89 | 20.73 |
| Ligands | 19.14 | 18.95 | 19.18 | 20.60 | 19.09 |
| Water | 40.35 | 37.10 | 40.75 | 46.94 | 40.71 |
| Clashscore | 2.20 | 1.89 | 2.20 | 2.20 | 1.57 |
| Ramachandran |  |  |  |  |  |
| Favored (%) | 99.35 | 99.35 | 99.35 | 99.35 | 99.35 |
| Allowed (%) | 0.65 | 0.65 | 0.65 | 0.65 | 0.65 |
| Outliers (%) | 0.00 | 0.00 | 0.00 | 0.00 | 0.00 |

<sup>1</sup> Reported by *dials.scale* in *DIALS* [44]<sup>2</sup> Reported by *PHENIX* [50]

**Table S10.** Data reduction statistics for DHFR EF-X from Laue diffraction

| Dataset | Off | Off (reduced sym.) | 200 ns (+3.5 kV) | 200 ns (-3.5 kV) |
| --- | --- | --- | --- | --- |
| No. of Images | 363 | 363 | 363 | 363 |
| Spacegroup | $P2_12_12_1$ | $P2_1$ | $P2_1$ | $P2_1$ |
| Cell dim. (Å) |  |  |  |  |
| a | 34.29 | 34.29 | 34.29 | 34.29 |
| b | 45.53 | 45.53 | 45.53 | 45.53 |
| c | 99.00 | 99.00 | 99.00 | 99.00 |
| Total obs. | 723,372 | 723,372 | 710,019 | 709,472 |
| Unique obs. | 17,637 | 33,671 | 33,671 | 33,669 |
| Resolution (Å) | 41.36 - 1.70 | 41.36 - 1.70 | 41.36 - 1.70 | 49.50 - 1.70 |
|  | (1.76 - 1.70) | (1.76 - 1.70) | (1.76 - 1.70) | (1.76 - 1.70) |
| Multiplicity | 35.53 (27.40) | 18.63 (14.16) | 18.35 (13.73) | 18.34 (13.68) |
| Completeness (%) | 99.5 (99.4) | 99.4 (99.5) | 99.4 (99.5) | 99.4 (99.5) |
| Mean $F/\sigma_F$<sup>1</sup> | 39.38 (19.51) | 28.54 (14.00) | 28.85 (13.86) | 28.85 (13.83) |
| CC<sub>1/2</sub><sup>1</sup> | 0.991 (0.957) | 0.987 (0.927) | 0.989 (0.929) | 0.988 (0.929) |

<sup>1</sup> Statistics were computed based on output from *careless* [46]

**Table S11.**
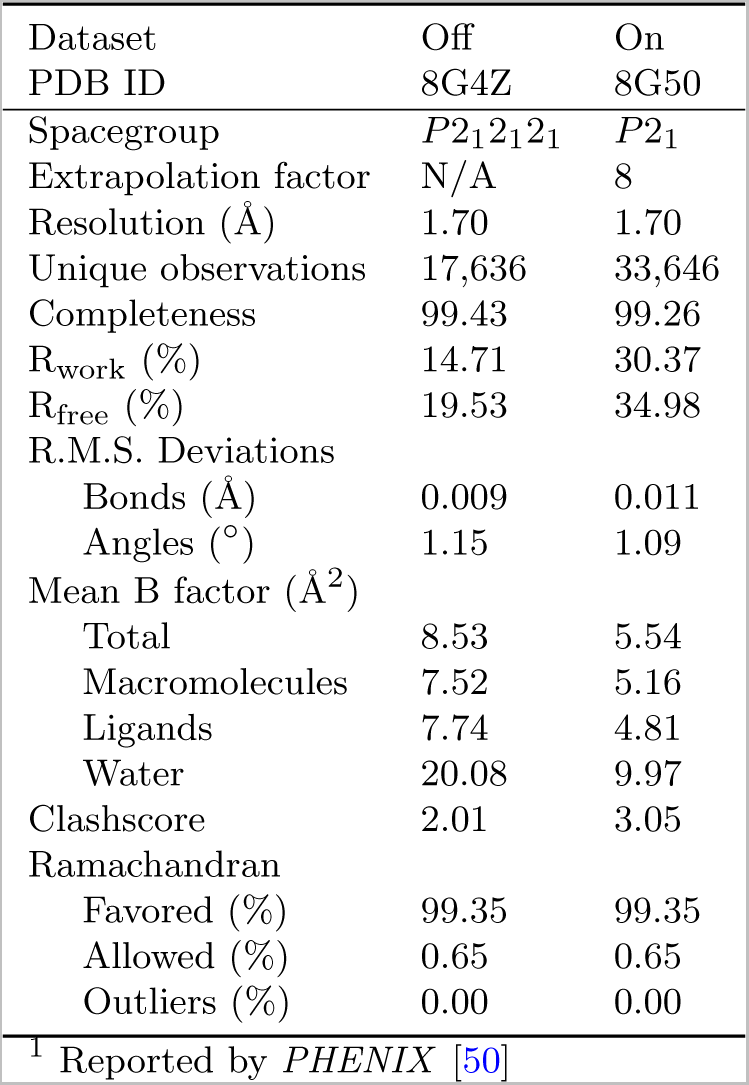
Refinement statistics for DHFR EF-X^1^

| Dataset | Off | On |
| --- | --- | --- |
| PDB ID | 8G4Z | 8G50 |
| Spacegroup | $P2_12_12_1$ | $P2_1$ |
| Extrapolation factor | N/A | 8 |
| Resolution (Å) | 1.70 | 1.70 |
| Unique observations | 17,636 | 33,646 |
| Completeness | 99.43 | 99.26 |
| R<sub>work</sub> (%) | 14.71 | 30.37 |
| R<sub>free</sub> (%) | 19.53 | 34.98 |
| R.M.S. Deviations |  |  |
| Bonds (Å) | 0.009 | 0.011 |
| Angles (°) | 1.15 | 1.09 |
| Mean B factor (Å<sup>2</sup>) |  |  |
| Total | 8.53 | 5.54 |
| Macromolecules | 7.52 | 5.16 |
| Ligands | 7.74 | 4.81 |
| Water | 20.08 | 9.97 |
| Clashscore | 2.01 | 3.05 |
| Ramachandran |  |  |
| Favored (%) | 99.35 | 99.35 |
| Allowed (%) | 0.65 | 0.65 |
| Outliers (%) | 0.00 | 0.00 |

<sup>1</sup> Reported by *PHENIX* [50]

